# Ire1α-Regulated mRNA Translation Rate Controls the Identity and Polarity of Upper Layer Cortical Neurons

**DOI:** 10.1101/2021.06.23.449563

**Authors:** Mateusz C. Ambrozkiewicz, Ekaterina Borisova, Andrew G. Newman, Matthew L. Kraushar, Theres Schaub, Rike Dannenberg, Marisa Brockmann, Marta Rosário, Paul Turko, Olaf Jahn, David R. Kaplan, Takao Iwawaki, Christian M. T. Spahn, Christian Rosenmund, Victor Tarabykin

## Abstract

Evolutionary expansion of the neocortex is associated with the increase in upper layer neurons. Here, we present Inositol-Requiring Enzyme 1α, Ire1α, as an essential determinant of upper layer fate, neuronal polarization and cortical lamination. We demonstrate a non-canonical function of Ire1α in the regulation of global translation rates in the developing neocortex through its dynamic interaction with the ribosome and regulation of eIF4A1 and eEF-2 expression. Inactivation of Ire1α engenders lower protein synthesis rates associated with stalled ribosomes and decreased number of translation start sites. We show unique sensitivity of upper layer fate to translation rates. Whereas eEF-2 is required for cortical lamination, eIF4A1 regulates acquisition of upper layer fate downstream of Ire1α in a mechanism of translational control dependent on 5’UTR-embedded structural elements in fate determinant genes. Our data unveil developmental regulation of ribosome dynamics as post-transcriptional mechanisms orchestrating neuronal diversity establishment and assembly of cortical layers.

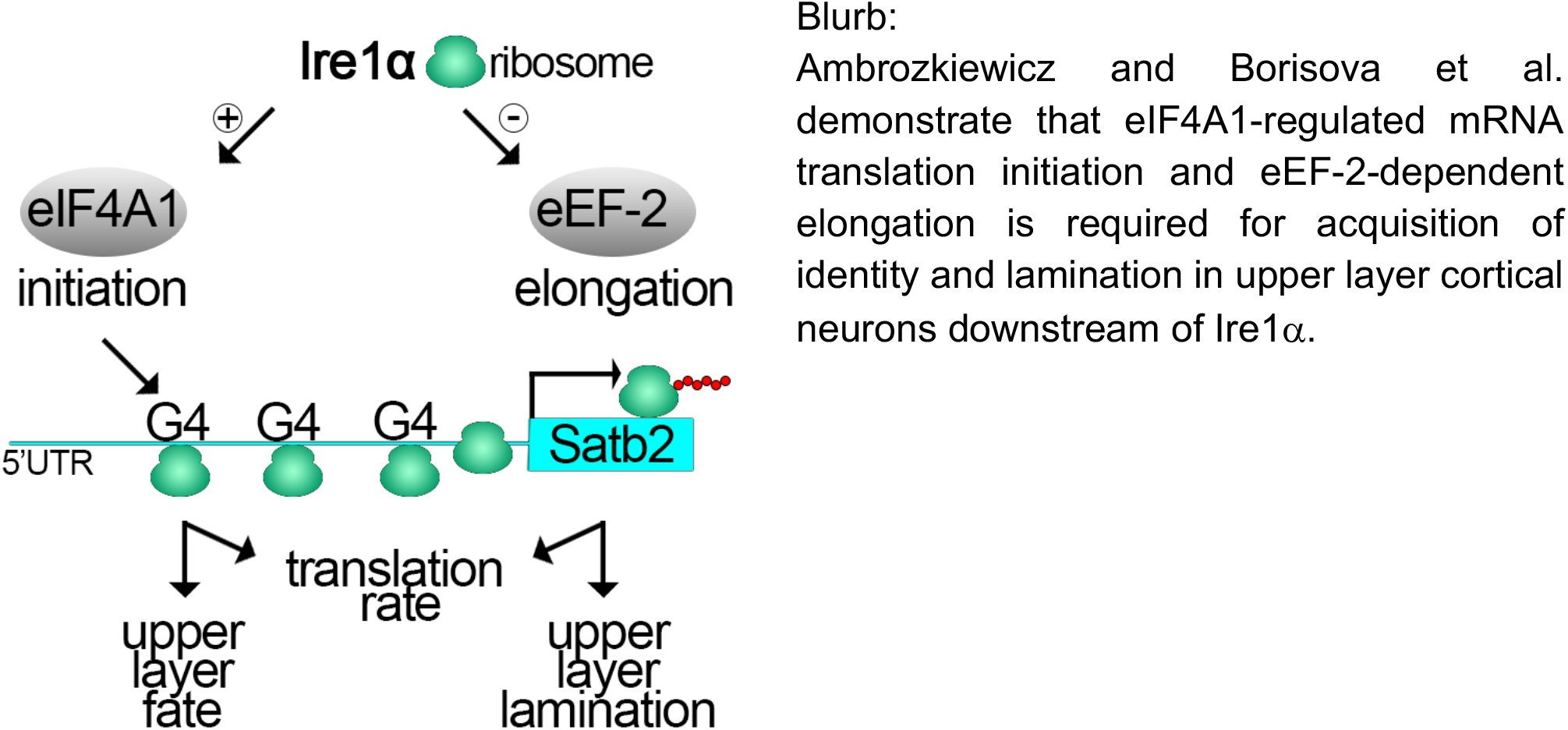

**HIGHLIGHTS:**

- Small molecule screening reveals Ire1α upstream of upper layer neuronal identity
- Polarization and proper lamination of layer II/III neurons require Ire1α
- Development of upper layers requires high translation rates driven by eIF4A1 and eEF-2 downstream of Ire1α
- eIF4A1-dependent Satb2 mRNA translation initiation is a mechanism of upper layer fate acquisition

## INTRODUCTION

The provenance of cellular diversity in the cerebral cortex has been the center of attention for developmental neuroscientists for decades. The mammalian neocortex, evolutionarily the newest add-on to the central nervous system, is organized in six laminae, each populated by neurons with distinct afferents and efferent connectivity patterns, transcriptomic signatures, and electrophysiological properties. Pyramidal glutamatergic neurons of the neocortex are born from the progenitors at the ventricular side of the brain primordium and polarize synchronously with their migration to laminate distinct cortical layers. Deeper layers, forming subcortical tracts, are established first. Later born, upper layer neurons migrate through pre-established deeper layers, and differentiate to extend polarized axons forming intracortical tracts, such as the corpus callosum (*1–6*).

Transcription factor signatures characterize pyramidal neurons of different layers. Special AT-Rich Sequence-Binding Protein 2 (Satb2) is one of the strongest determinant of intracortically projecting pyramidal neurons identity and is required to from the corpus callosum (*7, 8*). Conversely, subcortical projections are determined by expression of FEZ Family Zinc Finger (Fezf) and COUP-TF-Interacting Protein 2 (CTIP2, also known as Bcl11b) in deep layer V and VI neurons, where loss of either gene disrupts the corticospinal tract (*9*–*12*).

The temporal progression of ventricular progenitor state requires restriction of multipotency towards generating solely upper layer Satb2-positive neurons. Current models describe the temporal progression of neuronal differentiation as conserved across deep and upper layers, but the initial ground states embedded in their progenitors as progressively discrete (*13–15*). Transcriptomic signatures in apical progenitors are temporally plastic, and able to adapt to the developmental milieu in elegant heterochronic transplantation experiments (*16*). Transcriptional priming in progenitors involves translational repression which restricts expression of early lineage genes in the later born lineage (*17*).

The models described above focus heavily on transcriptional measurements. In addition to these, we and others have previously demonstrated that post-transcription, including RNA-binding proteins, translation, and ubiquitination pathways are major regulators of cortical connectivity, neuronal subtype specification, and axonal projections (*18, 19*). In the mature brain, high-fidelity protein translation is proven essential to maintain synaptic transmission (*20, 21*). However, how key developmental events are regulated at the level of translation, including neuronal polarization and the temporal succession of cortical progenitor fate, remains unclear.

In this study, we sought upstream regulators of Satb2, which determines the specification of callosally projecting cortical neurons (*7, 8*). Using a genetic reporter, we screened a library of small molecules for their ability to alter Satb2 expression in developing neurons. Among our most impactful candidates was an inhibitor of kinase/RNase Inositol-Requiring Enzyme 1α (Ire1α), also known as ER-to-Nucleus Signaling 1 (Ern1), the main sensor of ER lumen homeostasis and regulator of the Unfolded Protein Response (UPR). Upon ER stress, a translational shift promotes expression of proteins vital for cell survival and restoration of the ER folding capacity (*22*). Additionally, Ire1α regulates stress-independent remodeling of actin filaments by its association with filamin A (*23*), linked to periventricular heterotopia (*24*). Another study unveiled a direct interaction of Ire1α with the ribosomes *in vitro* (*25*). Beyond that, homeostatic functions of Ire1α, especially in the developing neocortex, have remained quite elusive.

In this work, we demonstrate that Ire1α is essential for acquisition of Satb2 neuronal identity, neuronal polarization and lamination during corticogenesis. Conditional deletion of *Ire1α* in the neocortex results in global decrease of translation rates, lower number of translation sites associated with decreased level of of eukaryotic initiation factor 4A1 (eIF4A1) and slower elongating ribosomes. Moreover, we reveal that the development of polarized upper layer neurons requires high rates of protein translation. We provide evidence that upper layer fate acquisition requires Ire1α-regulated and 5’UTR-embedded eIF4A1-dependent translational control of Satb2 itself. Taken together, our study defines translational requirements of neuronal progenitors and distinct neuronal subtypes, unveiling the layers of gene expression regulation driving neuronal diversity during corticogenesis. Additionally, this study extends the function of Ire1α beyond stress pathways to an innate developmental signal for neuronal specification in the cortex.

## RESULTS

### Ire1α is a positive regulator of Satb2 and axon specification in developing cortical neurons

In order to identify signaling pathways orchestrating the balance between upper and deeper layer neurons in the cortex, we have previously developed a method for investigating cell fate acquisition (*26*), which utilizes the *Satb2*^Cre/+^ mouse line (*27*). By expressing a Cre-inducible fluorescent reporter loxP-Stop-loxP-tdTomato in E13.5 *Satb2*^Cre/+^ primary cortical cells (Fig. 1a), we used cell sorting to quantify the proportion of Satb2-expressing tdTomato-positive cells (*Satb2*^tdTom^). Co-transfection with EGFP-encoding plasmid allowed to normalize for the transfection efficiency.

**Fig. 1.**
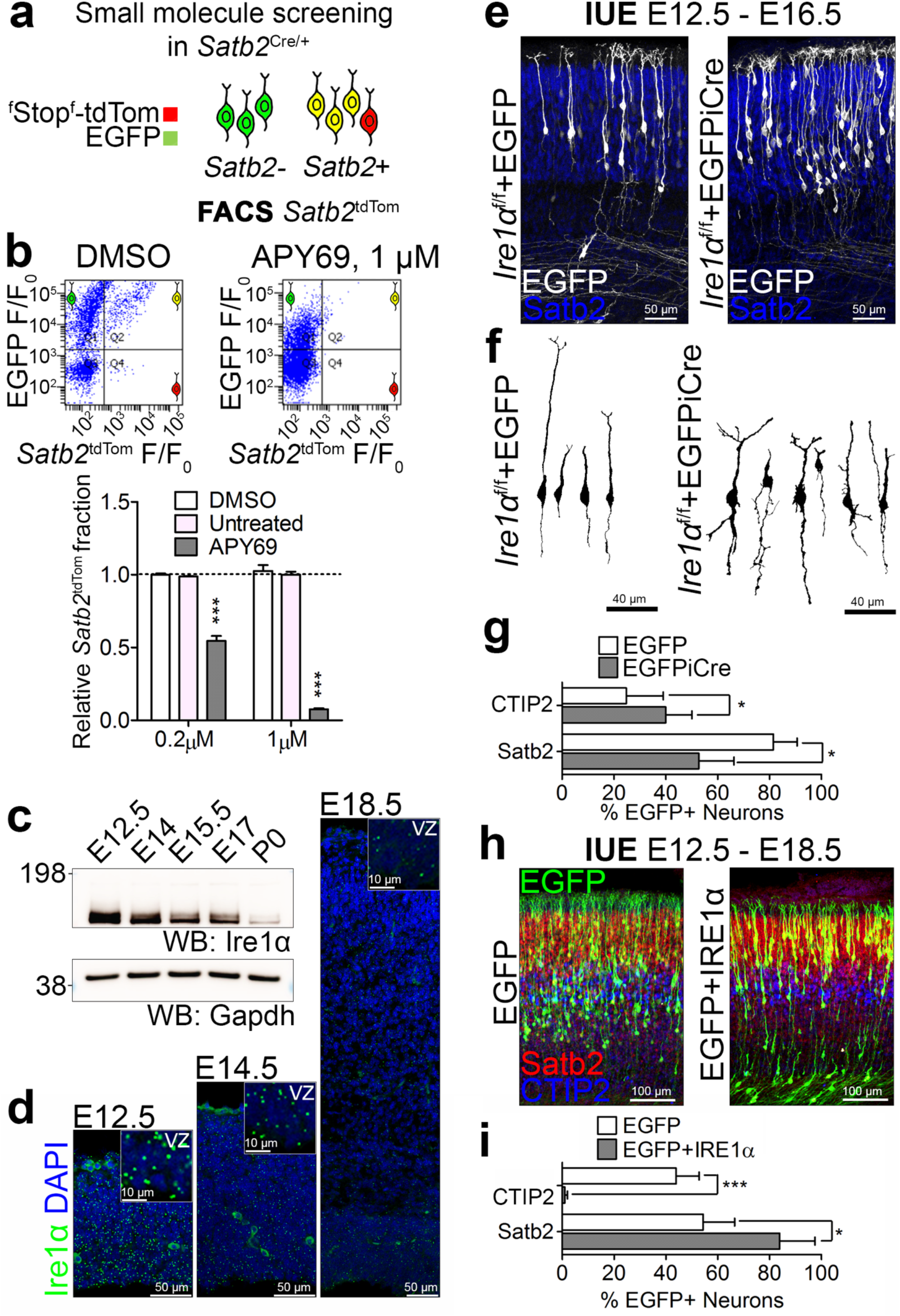
Small molecule screening reveals Ire1α activity pivotal for upper layer neuronal identity and bipolar morphology of cortical neurons. (a) Screening workflow to identify signaling pathways upstream of Satb2 neuronal identity. (b) Top panels: representative results of flow cytometry of DIV2 neurons treated with DMSO and APY69, a selective inhibitor of Ire1α. Bottom panels: quantification of the proportion of *Satb2*-positive neurons. (c) Western blotting in cortical lysates to profile developmental expression of Ire1α. Gapdh, loading control. (d) Fluorescence in situ hybridization for Ire1α in cortical sections at indicated developmental stages. Insets demonstrate enlarged fragments of the ventricular zone (VZ). (e) Representative images of immunostaining against EGFP and Satb2 in E16.5 coronal cortical sections of *Ire1α*^f/f^ embryos after IUE at E12.5 with plasmids encoding for EGFP or EGFP and Cre simultaneously. (f) Example EGFP-based tracings of single neurons after IUE in (e). (g) Quantification of neuronal cell identity after IUE described in (e). (h) Representative images of cortical coronal sections at E18.5 after IUE in wild-type E12.5 embryos to express EGFP, or EGFP and human IRE1α. (i) Quantification of neuronal cell identity after IUE described in (h). For (e), (g) and (h-i), compare Fig. S7. Bar graphs show averages ± S.D. Statistics for (b), one-way ANOVA with Bonferroni post-hoc test; (g) and (i), D’Agostino-Pearson normality test and unpaired t-test. *** p < 0.001; 0.01 < * p < 0.05.

Using this system, we screened a library of small molecule inhibitors for their ability to alter the proportion of *Satb2*^tdTom^ neurons at day-in-vitro two (DIV2) (*28*). APY69 was among the strongest modulators of Satb2 expression with its dose-dependent effect (Fig. 1b, Table S1). APY69 specifically inhibits Ire1α, an evolutionarily ancient bimodal transmembrane kinase and RNase, which acts as an endoplasmic reticulum (ER) stress sensor (*29, 30*). Recent evidence highlights an unfolded protein response (UPR)-independent developmental function of Ire1α as regulator of cellular migration (*23*). We hypothesized that stress-independent functions of Ire1α might be essential for specification of Satb2 identity in developing neurons.

We first studied the expression pattern of Ire1α. Fluorescence in situ hybridization (FISH) and Western blotting revealed a strong developmental downregulation of Ire1α, expressed homogenously throughout the developing cortex, with its highest levels at E12.5, both at the level of protein, and mRNA (Fig. 1c-1d)

We then went on to validate the pharmacological screening results for APY69 (Fig. 1b) using a knock-out (KO) mouse model. Given ubiquitous expression of Ire1α in many tissue types, its full KO results in overt developmental defects, associated with non-central nervous system-related phenotypes (*31*). To carve out the role of Ire1α in cortical development, we used the *Ire1α*^f/f^ line, which enables Cre-dependent deletion of exons 20–21 from the floxed *Ire1α* allele. Upon expression of Cre, the kinase-extension nuclease (KEN) domain including RNase active site is deleted from the Ire1α protein.

To inactivate *Ire1α* in defined cell populations in the developing cortex, we took advantage of *in utero* electroporation (IUE), which allows for *in vivo* DNA delivery to a spatiotemporally defined subset of progenitors at the ventricular surface of the embryonic cortex. Early (E12.5) neuronal progenitors display multipotency in their ability to generate both CTIP2-and Satb2-positive neuronal progeny, whereas late (E14.5) progenitors give rise to exclusively Satb2-expressing neurons populating upper layers. In our first set of experiments, we used IUE in *Ire1α*^f/f^ mice to transfect EGFP as a control and both EGFP and Cre to induce *Ire1α* KO in a subset of early cortical progenitors at E12.5 and derived progeny (Fig. 1e). We then immunostained E16.5 coronal cortical sections for EGFP and neuronal identity markers, Satb2 and CTIP2. We noted disrupted, highly branched morphology of *Ire1α* KO neurons, compared to majority of bipolar control cells (Fig. 1f). Quantification revealed fewer Satb2-expressing neurons at the expense of CTIP2-positive ones in the *Ire1α* KO (Fig. 1g), concordant with our *in vitro* results (Fig. 1b). On the contrary, overexpression of human IRE1α in wild-type early cortical progenitors at E12.5 (Fig. 1h) resulted in an increased proportion of Satb2-expressing neurons at E18.5, and remarkably, hardly any CTIP2-positive neurons (Fig. 1i).

Post-mitotic promoter NeuroD1-induced expression of Cre in *Ire1α*^f/f^ at E12.5 did not alter the proportion of Satb2 and CTIP2 neurons at E16.5 (Fig. S1a and S1b), indicative of a neuronal progenitor-embedded role of *Ire1α* in the acquisition of neuronal identity. We observed an analogous morphological phenotype in NeuroD1-Cre expressing neurons in *Ire1α*^f/f^ mice as in ones transfected with CAG-Cre (Fig. S1a and S1c, compare with Fig. 1f).

Using an Emx1-Cre deleter, we then conditionally inactivated *Ire1α* in the progenitors of dorsal telencephalon by establishing a *Ire1α*^f/f^; *Emx1*^Cre/+^ mouse line, *Ire1α* cKO. We injected pregnant mice with BrdU at E12.5 (Fig. S2a-S2d) and E14.5 (Fig. S2e-S2h) and quantified the proportions of Satb2 and CTIP2-expressing neurons at P2, when neurogenesis is complete. Among all E12.5-derived BrdU+ cells, we counted fewer Satb2+ and more CTIP2+ cells (Fig. S2b and S2c) in the cKO, consistent with our IUE experiments using *Ire1α*^f/f^ line (Fig. 1). Moreover, in E14.5 BrdU-pulsed P2 cKO cortices, we found more BrdU+/CTIP2+ cells than in control cortices which localized to the top of the CP (Fig. S2f and S2g).

These data altogether indicate a critical requirement of Ire1α for the correct proportion between upper layer-associated Satb2-positive and deeper layer-associated CTIP2-expressing neurons specified during cortical development.

### Ire1α is required for axon specification in developing cortical neurons of upper layers

Efferent cortical connectivity is established by the correct proportion of neuronal types and specification of functional axons during development. Because of the prominent morphological phenotype for E12.5 progenitor-derived neurons (Fig. 1f), and given that pyramidal upper layer neurons serve as an established model to study specification of axon-dendrite polarity (*32*), we electroporated E14.5 neuronal progenitors of *Ire1α*^f/f^ embryos with plasmids encoding EGFP or EGFP and Cre and fixed the brains at E18.5. In line with previously published results (*23*), loss of *Ire1α* lead to a disrupted laminar positioning of upper layer cortical neurons with a substantial number of Cre-expressing cells populating the bottom of the cortical plate (CP; Fig. 2a and 2b). Consistently, in the cKO brains, both E12.5 and E14.5 BrdU-pulsed Satb2-expressing cells localized to the lower portions of the CP, as compared to the control cells (Fig. S2). Using expansion microscopy (ExM), we detected multiple trailing processes (TPs) and many short processes originating from the soma in *Ire1α* KO upper layer neurons (Fig. 2c). This contrasted with the majority of control neurons characterized with a single TP and lack of additional neurites emanating from the soma at this stage (Fig. 2d). Disruption of bipolar morphology and aberrant laminar positioning within the CP was independent of Satb2 expression (Fig. 2e). We also observed morphological defects induced by IUE of NeuroD1-Cre inducing *Ire1α* KO in differentiating neurons (Fig. S1d-S1f). These data indicate Ire1α has both a progenitor and post-mitotic neuron embedded role in the regulation of neuronal morphology during cortex development.

**Fig. 2.**
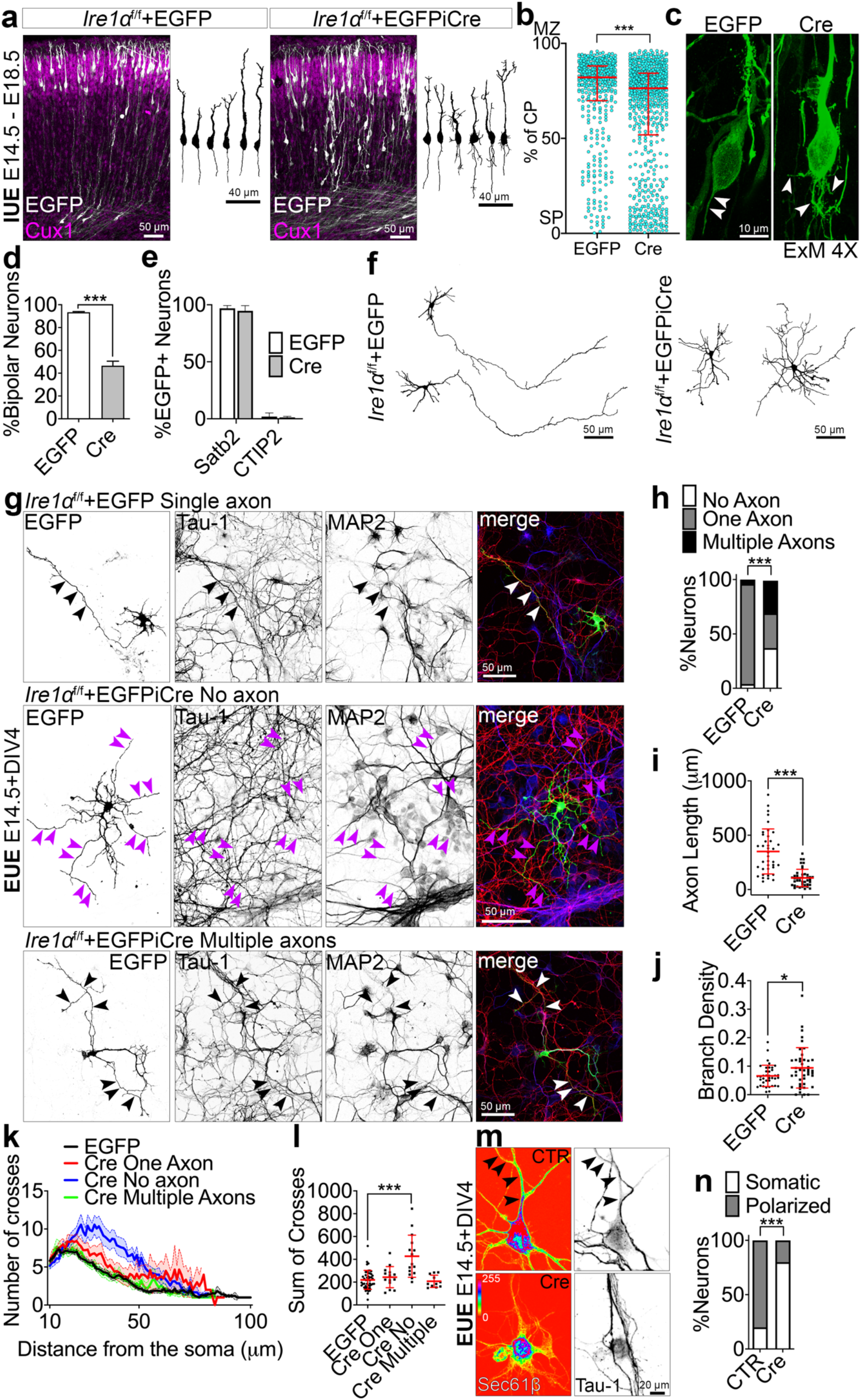
Ire1α is indispensable for bipolar morphology, axon specification and proper laminar positioning of upper layer neurons. (a) Representative images of EGFP and upper layer-expressed Cux1 immunostaining in coronal cortical sections from e18.5 *Ire1α*^f/f^ embryos after IUE at e14.5 with indicated plasmids and semi-automatic, EGFP-fluorescence based tracings of single neurons. (b) Laminar distribution of cortical neurons in e18.5 *Ire1α*^f/f^ brains after IUE described in (a). The position of each neuron was normalized to the thickness of the CP and represented as a dot. 0% - Subplate (SP), 100% - Marginal Zone (MZ). Graph contains pooled data from indicated number of brains (Table S1). (c) Representative images of EGFP immunostaining after expansion microscopy (ExM) in cortical sections from (a). Arrowheads point to trailing processes. (d) Average proportion of bipolar neurons from (a). (e) Quantification of neuronal identities in experiment in (a). (f) Representative EGFP-based tracings of *Ire1α*^f/f^ neurons at DIV4 after EUE to express EGFP or EGFP and Cre. (g) Neurons in (f) were fixed and immunostained with axonal or dendritic markers. White and black arrowheads indicate axons, purple ones indicate neurites lacking Tau-1 expression. (h) The number of axons projected from a single neuron. Average axon length (i), average axon branch density per micrometer (j), Sholl analysis diagram (k) and average sum of crossing dendrites (l) in neurons expressing EGFP or EGFP and Cre, projecting no, one, or multiple axons. (m) Representative EGFP signals and Tau-1 immunostaining of *Ire1α*^f/f^ DIV4 neurons after EUE to co-express EGFP-Sec61β and empty vector (CTR) or Cre. (n) Quantification of EGFP-Sec61β localization in experiment in (m). Line and error bars on (b) indicate median and interquartile range, scatter plots in (i), (j), (l) and bar graphs on (d) and (e) show averages ± S.D. Results on (k) are represented as averages ± S.E.M. Statistics for (b), (d-e), (i-j), D’Agostino-Pearson normality test and Mann-Whitney or unpaired t-test; (h), Chi-square test; (l), Kruskal-Wallis with Dunn’s multiple comparisons test; (n), Fisher’s test. *** p < 0.001; 0.01 < * p < 0.05.

To study the identity of excessive processes specified by *Ire1α* KO neurons, we expressed EGFP or simultaneously EGFP and Cre in E14.5 upper layer neuron progenitors of *Ire1α*^f/f^ embryonic cortices using *ex utero* electroporation (EUE) and cultured isolated primary cells until DIV4, when axons are established in control neurons (*18*). As expected, the majority of control cells projected a single longest neurite, while *Ire1α* KO cells were symmetrical (Fig. 2f). We then immunostained the fixed cells for axonal and dendritic markers (Tau-1 and MAP2, respectively) and counted the number of axons projected by single neurons (Fig. 2g). We failed to identify an axon in a fraction of *Ire1α* KO neurons, and identified a significant proportion of neurons that projected multiple axons, as compared to polarized control cells (Fig. 2h). Where present, *Ire1α* KO axons were shorter (Fig. 2i) and more branched (Fig. 2j), as compared to control ones. *Ire1α* KO neurons with no axons exhibited increased dendritic branching, possibly implicating an axon-to-dendrite identity switch of single neurites. Axon possessing *Ire1α* KO neurons exhibited a normal dendritic tree, when compared to controls (Fig. 2k and 2l). This indicates formation of excess axons upon *Ire1α* KO, rather than a dendrite-to-axon transition in neurons with multiple axons. Morphological defects in the KO were associated with somatic distribution of ER tubules visualized by EGFP-Sec61β expression as compared to its polarized localization in neurites of control cells (Fig. 2m and 2n). This indicates a possible involvement of Ire1α in ER*-*driven neuronal polarization.

We then asked if such profound morphological defects have consequences on axon-related neurophysiology. At DIV6, axons of upper layer control neurons are enriched for Tau-1 (Fig. S3a), Ankyrin G (Fig. S3b), and voltage-gated Na^+^ channels (Na_v_s; Fig. S3c). In *Ire1α* KO neurons, we observed a perinuclear somatic localization of these axonal markers, indicative of their aberrant subcellular localization. Current clamp recordings from *Ire1α*^f/f^ autaptic hippocampal neurons infected with lentiviruses encoding for EGFP or Cre showed a higher number of action potentials (APs) generated upon lower current injections (Fig. S3d and S3e). This might be attributable to more axon initial segments (AIS) specified by *Ire1α* KO neurons and/or their altered molecular composition, which increases the chance of firing at lower currents. Importantly, overall membrane integrity and conductance of *Ire1α* KO neurons were unaltered (Fig. S3f and S3g). Altogether this evidence reveals that loss of bipolar morphology in developing *Ire1α* KO neurons is associated with defects in axon specification, subcellular mislocalization of axonal markers and increased current sensitivity.

### Ire1α drives upper layer specification and axon formation in the neocortex by controlling expression of translation initiation and elongation factors

During the UPR, robust cellular reprogramming is driven by Ire1α signaling network (*33, 34*). Ire1α activation is paralleled by suppression of general translation (*35*). We hypothesized that also during development, Ire1α orchestrates upper layer subtype and axonal polarization by influencing protein translation and thereby affecting expression of key developmental determinants. To test this, we analyzed the ribosome profile in forebrain-specific *Ire1α* cKO, and we purified actively translating ribosomes *ex vivo* (*36*). Sucrose density gradients revealed higher level of polysomes in *Ire1α* cKO cortices (Fig. 3a).

**Fig. 3.**
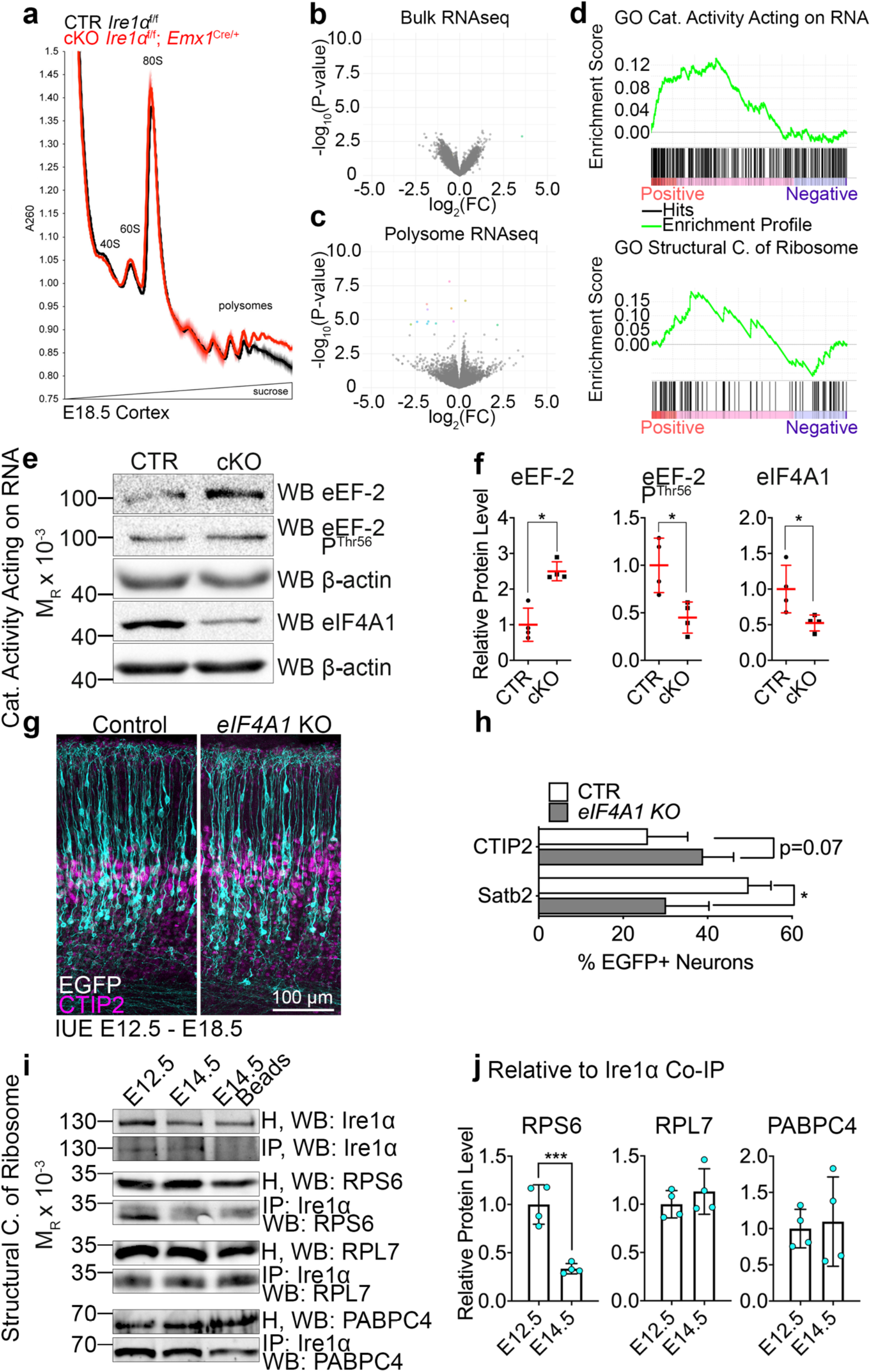
Ire1α-mediated protein translation initiation regulation drives upper layer identity acquisition in the cortex. (a) Analytic density gradient fractionation of A260-normalized E18.5 neocortex lysates, measuring the relative abundance of ribosomal subunits, 80S ribosomes, and polysomes. A260 curves plotted as mean ± S.D. across replicate fractionations (n = 2, each comprising 7-8 brains per condition), baseline (1.0) centered at onset of 40S peak. (b and c) Volcano plots for *Ire1α* cKO RNAseq in bulk tissue (b) and polysome-enriched RNA (c). Colored data points represent significantly regulated genes. FC; fold change. All P-values are Benjamini-Hochburg adjusted p-values. (d) Top two GSEA enrichment plots for cKO polysome RNA fraction versus control polysome RNA fraction, Catalytic Activity Acting on RNA and Structural Component of Ribosome (Table S2). (e) Representative Western blotting results using the control and *Ire1α* cKO E18.5 cortical lysates. (f) Quantifications of the protein level from (e). (g) Representative images of immunostaining against EGFP and CTIP2 in E18.5 coronal cortical sections of wild type embryos after IUE at E12.5 with gRNAs and Cas9 nickase to achieve indicated genotypes. (h) Quantification of neuronal cell identity in the experiment in (g). Compare Fig. S7. (i) Representative Western blotting results of endogenous Ire1α co-immunoprecipitation from E12.5 and E14.5 cortical lysate. Interaction between Ire1α and indicated proteins was quantified relative to the amount of immunoprecipitated Ire1α. Red line and error bars on (f) and bar graphs in (h) and (j) indicate mean ± S.D. Statistics for (f), (h), D’Agostino-Pearson normality test and Mann-Whitney test; for (j), Shapiro-Wilk normality test and unpaired t-test. *** p < 0.001; 0.01 < * p < 0.05.

We then investigated which RNAs localize to polysome fraction in *Ire1α* cKO cortex. We performed RNA sequencing in bulk cortex of the control and cKO as well as in the polysome fraction purified from the control and *Ire1α* cKO cortices. Notably, we did not detect gross changes in the level of the transcriptome in the bulk RNAseq [(Fig. 3b; cutoff at corrected p-value 0.05, AY036118 lncRNA outlier comes from inflated expression counts given its alignment with *Rn45s* repeat (*37*)]. Polysome-associated RNA in *Ire1α* cKO cortices (depicted as colored data points, cutoff at corrected p-value 0.05) encoded for structural components of the ribosome, proteins acting on RNA, revealed by the gene set enrichment analysis (GSEA, Fig. 3d), and other molecules regulating translation and initiation factors (Table S2). These results indicate that *Ire1α* loss in the cortex alters translation of RNAs crucial for ribosome complex function.

We then investigated the two pathways regarding their roles in cortical development. First, we quantified expression levels of canonical translation regulators (Fig. S4a and S4b) and identified upregulated levels of eukaryotic elongation factor 2 (eEF-2), downregulated phosphorylated eEF2 on Thr56 and decreased level of eukaryotic initiation factor 4A1 (eIF4A1) in the *Ire1α* cKO cortex (Fig. 3e and 3f). Given no gross changes on the RNA level (Fig. 3b), we conclude that the expression level changes for these factors are of translational nature. Notably, the level of spliced Xbp1, a cellular stress-associated Ire1α substrate was not altered, indicating a non-canonical UPR-independent role of Ire1α in the regulation of polysome level.

We then overexpressed eEF-2 using IUE in e12.5 and e14.5 wild type embryos (Fig. S4c-S4g). Four days later, we detected no alteration of Satb2 proportion in the neuronal progeny after eEF-2 OE (Fig. S4d and S4f). However, we did detect a positioning defect of upper layer neurons, similar to the one observed upon loss of *Ire1α* (Fig. S4g). Next, we inactivated *eIF4A1* using CRISPR-Cas9 technology (Fig. 3g), which allows for simultaneous sgRNA and humanized Cas9 nickase delivery to developing cortex using IUE in e12.5 wild type embryos (*38, 39*). The loss of *eIF4A1* resembled *Ire1α* KO (Fig. 1e and 1g) regarding the types of neurons generated, with less Satb2-expressing neurons, at the expense of CTIP2-positive cells (Fig. 3h). Loss of *eIF4A1* from upper layer progenitors did not affect laminar positioning in the cortex at e18.5 (Fig. S4e-S4g). Both eIF4A1 and eEF-2 were highly and homogenously expressed in the E12.5 cortex, and downregulated at E14.5, resembling the expression pattern of Ire1α (Fig. S4h-S4i, compare Fig. 1d). We also studied the polarity acquisition in DIV4 neurons after EUE-induced eEF-2 OE, *eIF4A1* KO and simultaneous eEF-2 OE and *eIF4A1* KO (Fig. S4j-S4k). We found that all three manipulations lead to emergence of neurons with 0 axons, characterized by absence of Tau-1 enrichment in a single neurite, partially reminiscent of *Ire1α* KO neurons (Fig. 2g-2h). Notably, simultaneous eEF-2 OE and *eIF4A1* KO resulted in the highest proportion of neurons without an axon.

Next, we hypothesized that the interaction between Ire1α and the ribosome is developmentally regulated. Using co-immunoprecipitation in E12.5 and E14.5 cortical homogenate, we quantified that endogenous Ire1α interaction with ribosomal protein S6, (RPS6), was significantly stronger at E12.5 (Fig. 3i and 3j). Binding to the components of the large ribosome subunit and interactors of mRNA poly(A) tail was not developmentally regulated. These results suggest a specific requirement for a strong interaction between Ire1α and small ribosomal subunit at the developmental timepoint for multipotent neuronal progenitors.

Altogether, these findings demonstrate that ribosome-associated Ire1α controls expression levels of translation regulators. In particular, initiation factor eIF4A1 acts downstream of *Ire1α* to control Satb2 expression and thereby neuronal subtype identity determination in the developing neocortex, while elongation factor eEF-2 regulates the laminar distribution of upper layer neurons. Tightly regulated levels of both eIF4A1 and eEF-2 in neuronal progenitors are critical for specification of an axon in developing neurons.

### Ire1α is a regulator of protein translation rates by controlling ribosome translocation and the number of translation sites

Increased polysome level in the *Ire1α* cKO may represent ribosomes elongating more slowly or stalling, and thus accumulating in the heavy fraction (*40*). To test this hypothesis, we examined translation rates in *Ire1α* cKO neurons using FUNCAT (*41, 42*). The method relies on replacing L-methionine with its alkyne analog L-homopropargylglycine (HPG) and detection in a Huisgen alkyne-azide cycloaddition reaction. The fluorescence intensity of azide-coupled Alexa-647 is proportional to the amount of incorporated HPG into newly synthesized proteins in defined time and serves as an estimate of translation rate in a given cell.

We first investigated protein synthesis rates in early E12.5 multipotent neuronal progenitors able to generate both upper and deeper layer neuronal lineages using EUE to induce *Ire1α* loss. At DIV1, we quantified lower translation rates in Cre-expressing mitotic marker Ki67-positive (*43*) *Ire1α* KO progenitors as compared to EGFP-expressing *Ire1α*^f/f^ ones (Fig. 4a). Similarly, we detected lower translation rates in E12.5 mitotic progenitors upon loss of *eIF4A1* (Fig. 4c and 4d). We quantified approximately 50% lower rates of translation in DIV4 upper layer *Ire1α* cKO neurons born from E14.5 cortical progenitors as compared to control cells (Fig. 4e and 4f). Taken together, regardless of the cell type, *Ire1α* loss engenders a decrease in the rate of protein synthesis. In the light of these data, *Ire1α* loss-mediated increase in polysome level may represent stalled or pausing ribosomes, accumulated in the polysome fraction.

**Fig. 4.**
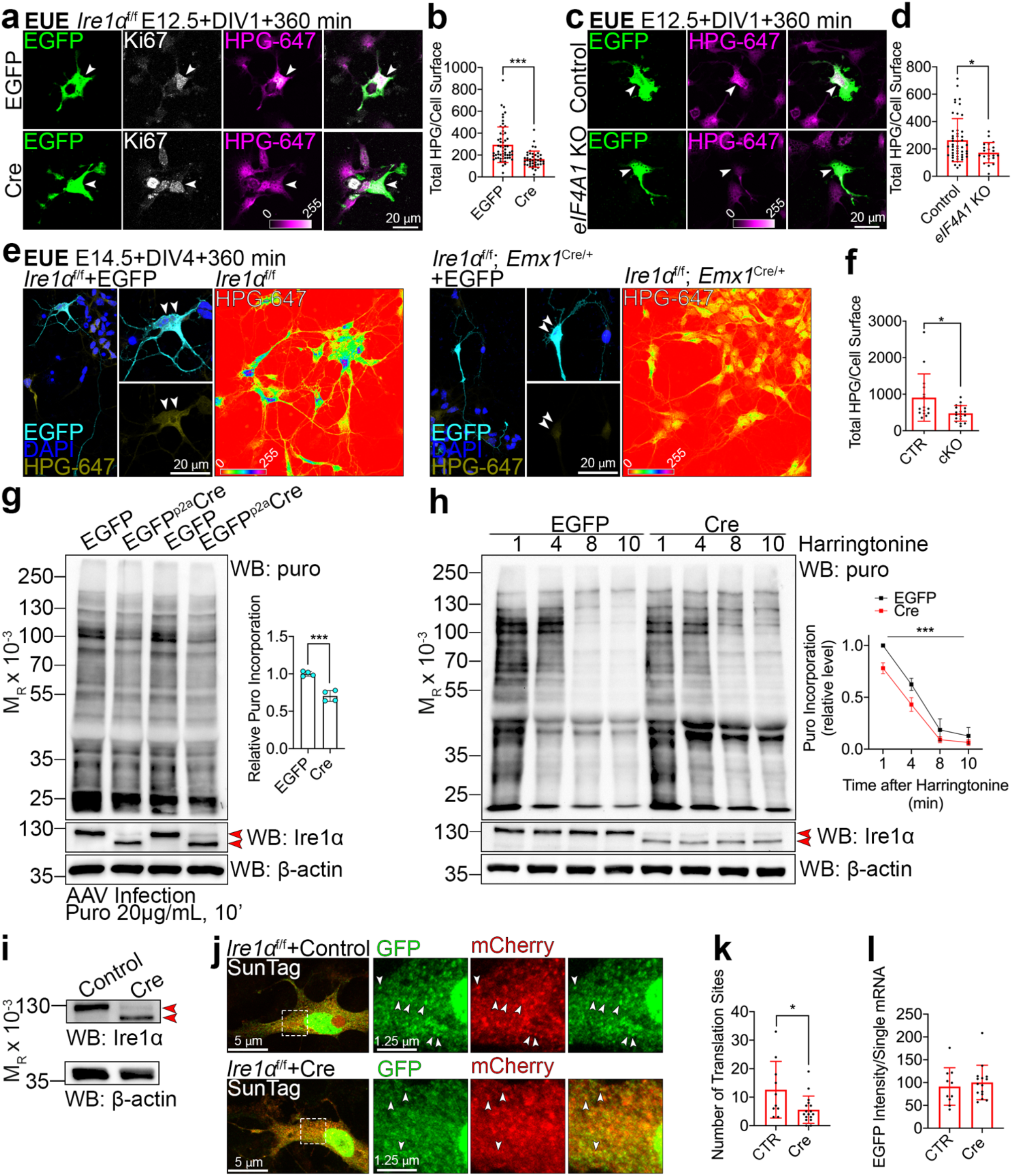
Loss of Ire1α results in suppression of translation rates as an effect of slower elongating ribosomes and decreased translation sites. (a), (c), (e) Images of representative primary cortical neurons prepared from *Ire1α*^f/f^ (a), wild type (c) or control and *Ire1α* cKO (e) embryos after EUE at E12.5 (a and c) or E14.5 (e) with indicated plasmids. Neurons were fed L-homopropargylglycine (HPG) for 360 min prior to fixation at DIV1 or DIV4. HPG was detected with Sulfo-Cyanine5 azide. (e) Right panels: images of HPG incorporation in control and cKO primary DIV4 neurons prepared from E14.5 cortex. White arrowheads point to Ki67-positive progenitors (a and c) or to somata of neurons derived from E14.5 progenitors (e). (b), (d), (f) Quantification of HPG incorporation. (g) Representative Western blotting using DIV5 lysates from *Ire1α*^f/f^ mouse embryonic fibroblasts (MEFs) after metabolic labeling of protein synthesis using puromycin and its quantification. MEFs were infected at DIV0 with control or Cre-expressing AAVs. Red arrowheads point to wild-type and KO form of Ire1α. (h) Representative Western blotting results of ribosome run-off assay using puromycin in control and KO MEFs at indicated timepoints after harringtonine treatment and quantification. (i) Western blotting validation of *Ire1α* KO in AAV-infected MEFs for the SunTag reporter experiment. (j) Representative images of empty and Cre-encoding virus infected MEFs expressing the SunTag24x-BFP-PP7 reporter. (k) Active translation sites were quantified in fixed MEFs. (l) Quantification of the intensity of scFv-GFP at translation sites. Bar graphs represent data points and averages ± S.D. Data point and error bars on (h) represent average ± S.E.M. from three independent experiments. Statistics for (b), (d), (f), (k), and (l) D’Agostino-Pearson normality test, Mann-Whitney, and for (l) unpaired t-test; (g), Shapiro-Wilk and unpaired t-test; (h), two-way ANOVA. 0.01 < * p < 0.05; *** p < 0.001.

To test this, we took advantage of ribosome run-off experiments with harringtonine (*44*). To circumvent a possibility of confounding effects of varying translation rates in different cell types, present in primary cortical cell culture models, we prepared murine embryonic fibroblasts (MEFs) from *Ire1α*^f/f^ embryos. We infected cells with Cre-encoding AAVs and kept in culture for DIV5 to ensure *Ire1α* KO. We first measured protein synthesis in MEFs using puromycin/antibody-based tool (*45*) and found that *Ire1α* KO MEFs showed diminished puromycin incorporation consistently indicative of reduced mRNA translation (Fig. 4g). *Ire1α* KO cells also showed diminished protein synthesis after the harringtonine treatment, reflecting slowly elongating ribosomes (Fig. 4h).

Next, we used SunTag (*46*) to label single mRNAs and nascent proteins in control and Cre-infected fixed *Ire1α* KO MEFs (Fig. 4i). To globally study translation dynamics in cells, mCherry-fused PP7 bacteriophage coat protein is used to identify mRNA and the EGFP-tagged single chain variable fragment recognizing the SunTag allows to detect nascent proteins. As compared to control, *Ire1α* KO MEFs demonstrated fewer colocalizing EGFP and mCherry puncta, reflective of decreased number of mRNA molecules in translation (Fig. 4j-4l). Altogether, we conclude that lower translation rates upon *Ire1α* KO are associated with slower ribosome elongation rates and decreased number of active translation sites.

### Deeper layer cortical neurons exhibit higher translation rates than upper layer neurons

Given lower translation rates in *Ire1α* cKO progenitors and neurons (Fig. 4), and our findings of Ire1α-mediated specification of neuronal subtype (Fig. 1), we next asked whether differences in translation rates are an intrinsic feature of postmitotic neurons of different cortical layers.

In order to test this, on the same day we nucleofected two populations of e12 and e14 primary cortical cells with dsRed and EGFP-expressing plasmids, respectively, mixed the now differentially labeled cells, and cultured together to tightly control for the microenvironment (to separate these sets of experiments from ones performed on *Ire1α,* we labeled the embryonic stages with a lowercase “e”). Our nucleofection and culture system enriches for deeper layer marker-expressing neurons derived from e12 cortex, for Satb2 in neurons prepared from e14 cortex, and generally for postmitotic neurons (Fig. 5a and 5b), with only a minor fraction of cycling cells or radial progenitors still present at DIV1 (Fig. 5c). We pulsed HPG in the methionine-free cell culture medium for 60, 120 and 240 minutes at DIV1 (for immature neurons) and at DIV5 (Fig. 4d). At DIV1, following HPG application, we quantified significantly higher HPG incorporation level in e12-(dsRed+) versus e14-(EGFP+) derived neurons (Fig. 5e and 5f). We also detected higher HPG incorporation rates at DIV5, with a milder difference between the two cell populations. Secondly, we observed, that in both e12-and e14-derived neuronal cultures, HPG incorporation rates were higher at DIV1 when compared to DIV5 (Fig. 5e and 5g). The addition of cycloheximide (CHX), a translation inhibitor (*47*), attenuates HPG incorporation (Fig. 5h), reinforcing the translation-specific nature of our findings. Altogether, these data indicate that deep layer-fated neurons have inherently higher translation rates compared to upper layer neurons.

**Fig. 5.**
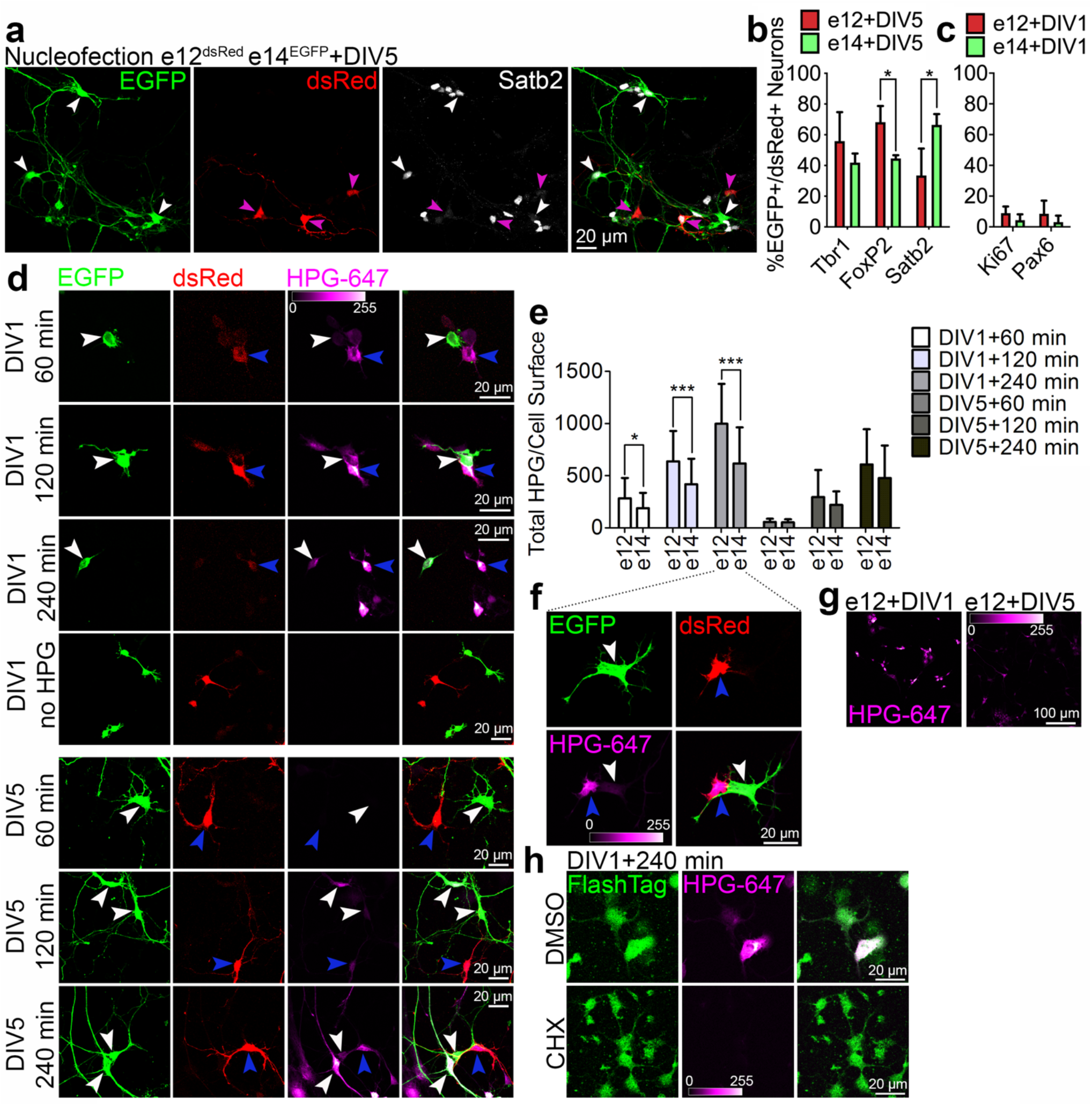
Increased translation rate is a hallmark of early born, deeper layer postmitotic neurons in the developing cortex. (a) Images of primary DIV5 neurons immunolabeled for EGFP, dsRed, and Satb2. Cortical cells from e12 embryo were nucleofected to express dsRed (purple arrowheads), and cortical cells from e14 embryo to express EGFP (white arrowheads). Both cell populations were mixed together and plated on a single glass coverslip. (b, c) Quantification of the cell identity in DIV5 neurons (b) and DIV1 primary cells (c) derived from e12 or e14 cortex. (d) Images of primary cortical cells immunolabeled for EGFP (white arrowheads) and dsRed (blue arrowheads), pulsed with HPG for indicated amount of time before fixation at DIV1 and DIV5. HPG was detected with far red fluorophore-coupled azide. (e) Incorporation of HPG was quantified as intensity of fluorescence signal normalized to the cell surface in primary neurons. (f) DIV1 primary cortical cells from (a), fed with HPG for 240 min. Note significantly higher HPG uptake in e12 cortex-derived immature neurons. (g) HPG labeling in primary neurons at DIV1 and DIV5 prepared from e12 cortex. Note significantly higher HPG uptake in primary cultures at DIV1. (h) DIV1 primary cortical cells from e14 cortex, fed with HPG for 240 min in presence of DMSO or 10 µg/mL cycloheximide (CHX). Note strongly reduced HPG labeling in cells cultured with CHX. FlashTag was added to the medium to label all cells. Bar graphs indicate mean ± S.D. Statistics for (b) and (c), D’Agostino-Pearson normality test and Mann-Whitney or unpaired t-test; (e), two-way ANOVA with Bonferroni post-tests. *** p < 0.001; 0.01 < * p < 0.05.

### The transition from multipotent neuronal progenitors to progenitors of upper layer neurons is associated with a pronounced upregulation of translation

Next, we studied the global translation rates in cycling progenitors of deeper and upper layer neurons and in their directly derived progeny. To address this, we used EUE, which unlike the nucleofection (Fig. 5) allows for labeling cycling progenitors. We transfected ventricular e12 progenitors with dsRed- and e14 progenitors with EGFP-encoding vectors. We then quantified HPG incorporation in Ki67-labeled DIV1 progenitors and in DIV5 postmitotic neurons (Fig. 6a) and found that e14 upper layer progenitors display profoundly higher translation rates as compared to e12 multipotent progenitors. Additionally, consistently with results shown in Fig. 5, e12 progenitor-derived postmitotic neurons display higher HPG incorporation rates as compared to e14 progenitor-derived cells. Translation rates undergo a significant upregulation in e14 cortical progenitors that dramatically decreases during postmitotic differentiation of an upper layer neuron (Fig. 6b). The majority of e12 progenitor-derived DIV5 postmitotic cells co-express both Satb2 and CTIP2 in culture, and e14 progenitor-derived ones express Satb2, but not CTIP2 (Fig. 6c and 6d).

**Fig. 6.**
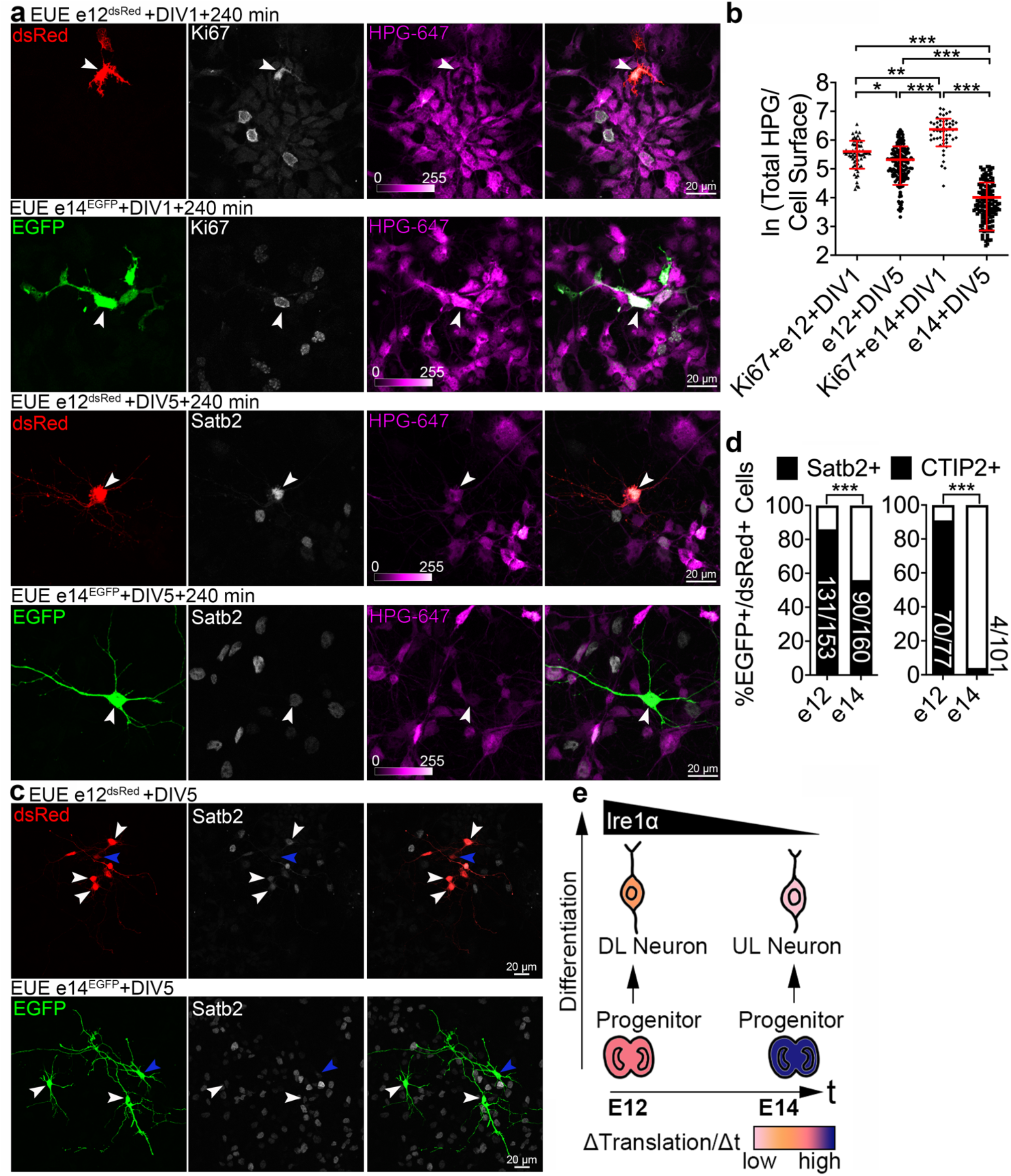
Progenitors of upper layers display dramatically higher translation rates than progenitors of deeper layers. (a) Images of primary DIV1 and DIV5 cortical cells immunolabeled for EGFP, dsRed, Ki67, and Satb2. To target Ki67-positive neuronal progenitors and their derived progeny, cortices of e12 embryos were *ex utero* electroporated (EUE) to express dsRed, and cortices of e14 embryos to express EGFP. Cells were triturated, mixed, plated together on a glass coverslip and pulsed with HPG for 240 min prior to fixation at DIV1 and DIV5. White arrowheads indicate representative cells. (b) Incorporation of HPG was quantified as intensity of fluorescence signal normalized to the cell surface area (natural logarithm scale). (c) e12 and e14-derived cells were fixed at DIV5 and immunolabeled for Satb2. White arrowheads point to neurons expressing high levels of Satb2, blue ones indicate example neurons with lower Satb2 expression. (d) Quantification of the cell identity markers in DIV5 neurons. (e) Schematic summarizing the translation rates (ΔTranslation/Δt) in different types of cortical cells for e12- and e14-derived lineage. The triangles indicate the dynamics of developmental enrichment for Ire1α; deeper layers (DL) and upper layers (UL). Graph on (b) indicates mean ± S.D. Statistics for (b), D’Agostino-Pearson normality test and Kruskal-Wallis test with Dunn’s multiple comparisons test; (d), Fisher’s test. *** p < 0.001; 0.001 < ** p < 0.01; 0.01 < * p < 0.05.

Taken together, the translation rate is a dynamic feature of cortical progenitors and their derived progeny, and likely represents cell- and stage-specific requirements of different protein sets during development.

### Transient attenuation of translation results in loss of axon and upper layer type identity

We then asked if pharmacologically attenuating translation early in development alters the specification of upper layer type neurons. Given a significant difference in translation rates at DIV1 between e12- and e14-derived neuronal cultures, we nucleofected the primary cells to express dsRed or EGFP as described above and exposed them to DMSO or cycloheximide (CHX), an inhibitor of protein synthesis, immediately after plating transiently for 24 hours. Following this, we changed the cell culture medium to one lacking CHX and maintained the primary neuronal cultures until DIV5 (Fig. 7a). Remarkably, transient inhibition of translation affected the appearance of Satb2 expression in cells derived from both stages but had no effect on the expression of CTIP2 (Fig. 7b).

**Fig. 7.**
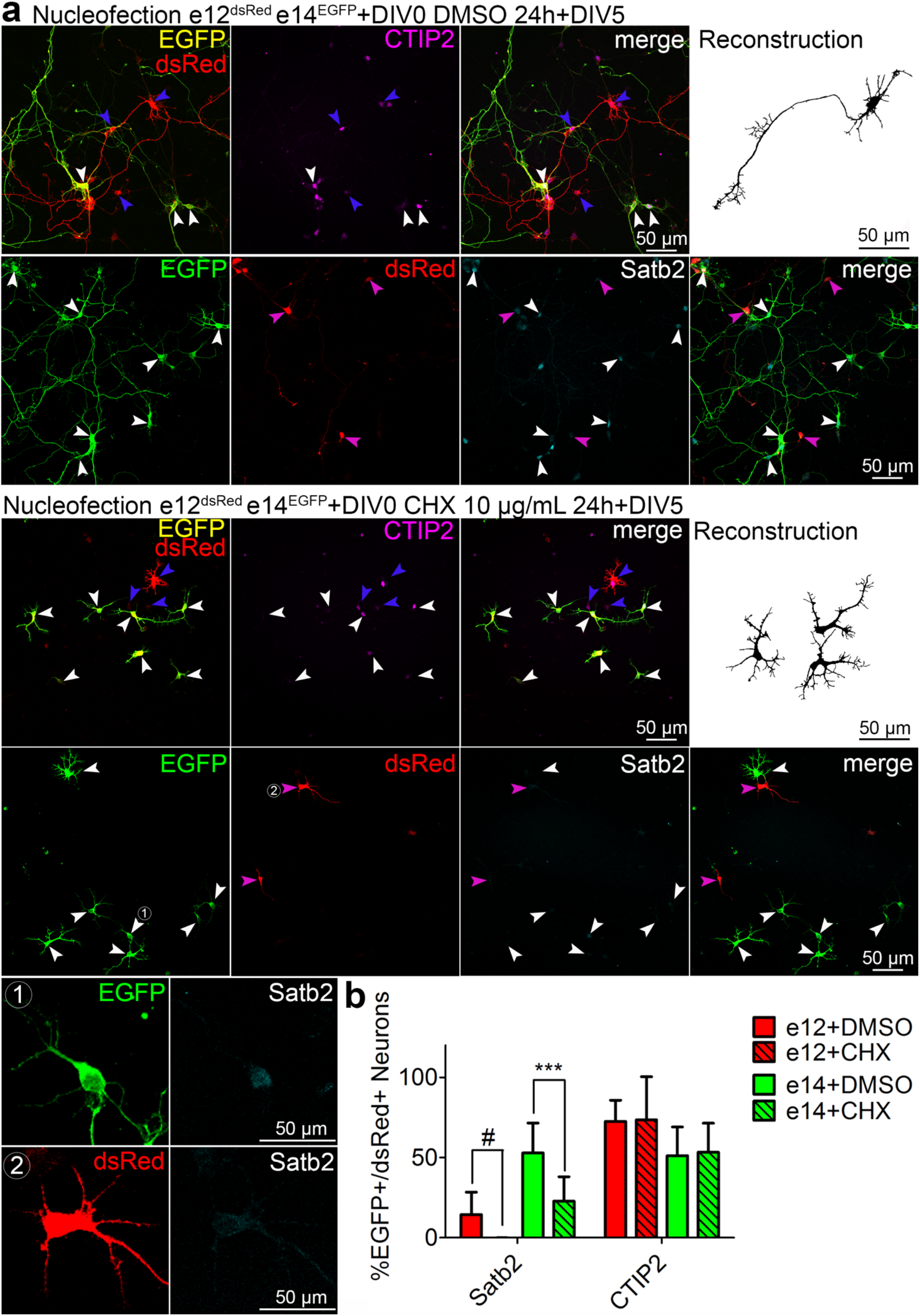
Upper layer identity requires a critical window of protein translation in precursor cells. (a) Images of immunolabeled primary cells from e12 embryo nucleofected to express dsRed, and from e14 embryo to express EGFP, mixed and plated on a single glass coverslip. Two hours post-plating, cells were treated with vehicle (DMSO) or cycloheximide (CHX) for 20 hours, followed by medium change, and fixed at DIV5. Upper panels show staining using rat anti-CTIP2, goat anti-EGFP, and rabbit anti-RFP, the latter one recognizing both EGFP and dsRed. For this reason, e12 cells in this case were recognized as solely expressing dsRed (blue arrowheads), but the e14 ones, both EGFP and dsRed (white arrowheads). Lower panels show anti-Satb2, anti-EGFP and anti-dsRed immunostaining with no cross-reacting antibodies; in this case, e12-derived cells express dsRed (blue arrowheads) and e14 ones EGFP (white arrowheads), as expected. Representative neuronal morphology is demonstrated as a semi-automated, EGFP- or dsRed-based reconstruction. (1-2) Example cycloheximide-treated e14 (1) or e12 (2) cortex-derived cells immunolabeled with an antibody anti-Satb2. (b) Quantification of the cell identity markers in DIV5 neurons derived from e12 or e14 cortex. Bar graphs indicate mean ± S.D. For statistics, two-way ANOVA with Bonferroni’s multiple comparisons test. # indicates a comparison between fractions of Satb2-positive e12 DMSO- and e12 CHX-treated group, the latter represented by virtually no positive cell. *** p < 0.001.

Additionally, neurons transiently exposed to CHX failed to project a long neurite at DIV5 (Fig. S5a). While control EGFP-expressing upper layer neurons displayed an asymmetric morphology with a longest neurite enriched for Tau-1 (Fig. S5a), most neurons transiently exposed to CHX failed to break their symmetry and expressed Tau-1 in the perinuclear regions of soma, resembling the scenario upon *Ire1α* (Fig. S3a, bottom panels). We quantified a CHX-induced dose-dependent loss of axons (Fig. S5b) and overall loss of neurite branching (Fig. S5c and S5d).

Altogether, transiently inhibiting protein synthesis at early developmental stages leads to irrevocable loss of Satb2 identity and loss of axons, indicative of critical translational window for both processes and reminiscent of *Ire1α* KO.

### Secondary structure of 5’UTR mRNA of Satb2 and CTIP2 regulates ribosome preference regulated by Ire1α and eIF4A1 during development

Given the critical requirement for high protein synthesis rates in neuronal progenitor for an upper layer fate and the regulation of global mRNA translation by Ire1α and eIF4A1, we investigated the mechanisms of Ire1α**-**mediated regulation of upper layer fate acquisition through direct Satb2 translation.

Translational control of eIF4A1 requires 5’UTR embedded elements, like G-quadruplexes (G4s) in the (especially) long and complex mRNA and allows for efficient scanning of translation start site (*48, 49*). We first modeled the 5’UTR structures of Satb2 and CTIP2 mRNA (Fig. 8a) and quantified predicted eIF4A1-dependent structural elements (Fig. 8b-8d). We found a higher density and the total number of G4s in 5’UTR of Satb2 as compared to CTIP2. We hypothesized that increased G4s in 5’UTR of Satb2 makes its mRNA translation uniquely sensitive to eIF4A1 and thereby also Ire1α level. Such regulation in primed progenitors displaying higher intrinsic translation rates would allow for acquisition of upper layer fate by the neuronal progeny.

**Fig. 8.**
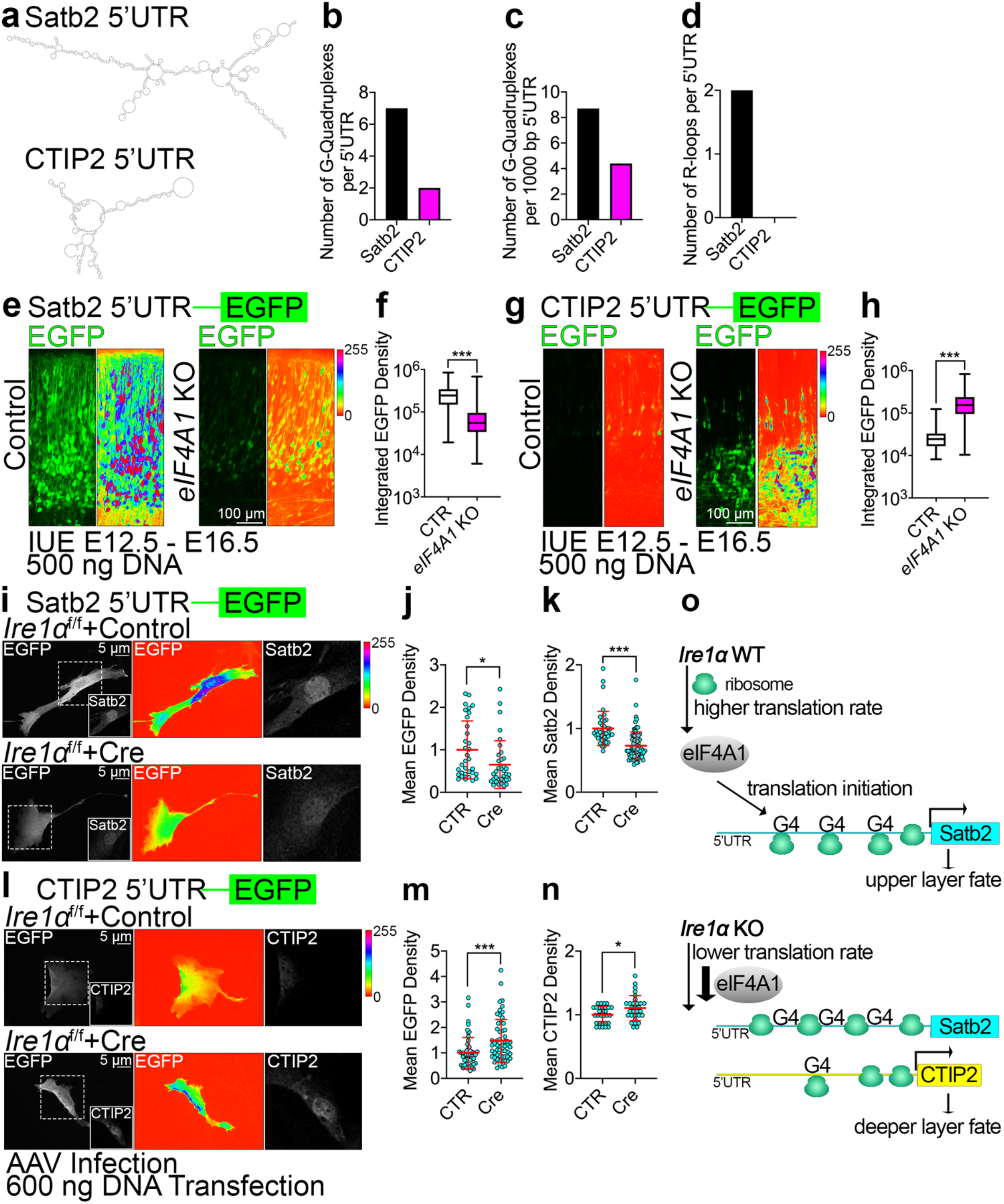
Helicase eIF4A1 and Ire1α regulate upper layer fate by controlling translation initiation through RNA secondary structural elements embedded in 5’UTR of Satb2. (a) Models of 5’UTR mRNA structures of Satb2 and CTIP2. (b-d) Quantification of putative G-quadruplexes and R-loops in the 5’UTR of Satb2 or CTIP2. (e, g) Representative images of EGFP fluorescence signals in the E16.5 brain sections after IUE at E12.5 with CRISPR-Cas9 vectors to achieve indicated genotypes and Satb2 5’UTR (e) or CTIP2 5’UTR (g) translation reporter construct. Shown are the native signals (left panels) and intensity encoding (right panels). (f, h) Quantification of EGFP fluorescence signals in single cells expressing translation reporters of Satb2 5’UTR (f) or CTIP2 (h). (i-n) Representative images of EGFP fluorescence signals (left panels: gray scale, middle panels: intensity encoding) and immunostaining for Satb2 (i) or CTIP2 (l) in *Ire1α*^f/f^ MEFs infected with Control or Cre-encoding AAVs at DIV0. At DIV5, infected MEFs were transfected with indicated reporter constructs, fixed and immunostained at DIV6. (j, m) Quantification of EGFP mean fluorescence signals of 5’UTR Satb2 reporter (j) or CTIP2 (m). (k, n) Quantification of mean nuclear fluorescence signals after immunostaining for Satb2 (k) or CTIP2 (n). (o) A summary of eIF4A1-dependent upper layer fate establishment, driven by Satb2 translation. Violin plots in (f) and (h) depict median, interquartile range (box) and minimum and maximum value (whiskers). Red line and error bars on (j-k), (m-n) indicate mean ± S.D. For statistical analyses, D’Agostino and Pearson normality test and Mann-Whitney test. *** p < 0.001; 0.01 < * p < 0.05.

We first constructed a fluorescence-based translational reporters by fusing 5’UTRs of Satb2 and CTIP2 to EGFP ORF. In these reporters, quantification of EGFP fluorescence intensity is a measure of 5’UTR-dependent translation efficiency. Next, we co-electroporated each reporter with CRISPR-Cas9 plasmids to induce *eIF4A1* KO in E12.5 cortical progenitors. We made sure the same amount of DNA was electroporated in control and KO condition. At E16.5, we observed a ribosome preference redistribution with dramatically decreased Satb2 reporter translation efficiency (Fig. 8e and 8f), and increased one for CTIP2 (Fig. 8g and 8h). This result demonstrates that translation efficiency of neuronal fate determinants depends on 5’UTR-embedded G4s and explains the mechanism of loss of upper layer fate in *eIF4A1* KO (Fig. 3g-3h).

Intriguingly, we observed the same tendencies for translational efficiencies for Satb2 and CTIP2 in *Ire1α* KO MEFs transfected with each reporter (Fig. 8i-8n). Additionaly, we demonstrated that lower translation efficiency of Satb2 5’UTR reporter is associated with decreased nuclear endogenous Satb2 expression (Fig. 8i, 8k). On the contrary, higher CTIP2 reporter translation was paralleled by increased nuclear CTIP2 expression in *Ire1α* KO MEFs (Fig. 8l, 8n).

We did not detect significant effect sizes using the reporters in *eIF4A1* KO and *Ire1α* KO at e14.5 (Fig. S6). These results indicate eIF4A1-driven timed regulation of translation downstream of Ire1α, embedded in 5’UTR of fate determinants, at early stages of corticogenesis, is required for generation of upper layer neurons.

## DISCUSSION

Development of the cortex comprises an orchestrated series of sequential, tightly controlled gene expression events with many layers of regulation. Our data implicate the regulation of protein synthesis rate by a mechanism involving eEF-2 and eIF4A1 downstream of Ire1α in the specification of upper layer neurons and their polarization. Through identifying a developmental role for Ire1α beyond its canonical role in translation stress pathways, we make the surprising finding that different protein synthesis rates are intrinsic features of distinct progenitor and differentiated neuron lineages.

The demand to swiftly and efficiently synthesize and remodel the proteome during cortical development places homeostatic pathways on the brink of cellular stress signaling. We speculate that such demands in normal development are met by the same molecular players as stress pathways, however, likely leading to unique downstream pathways active in development, such as a Satb2 program. During cortical development, Ire1α supervises the cellular translation flux (Fig. 4) by driving the expression of translation regulators and RNA binding proteins, essential for the high protein synthesis rate in upper layer progenitors, enabling translation of Satb2 in their postmitotic progeny (Fig. 3) and proteins crucial for specification of an axon. By labeling the amount of HPG incorporated into the newly synthesized proteins in a given time period, we quantify the total translation flux in a cell. HPG incorporation reflects several translation-regulating processes, e.g. the translation initiation, efficiency, polypeptide chain elongation, codon dwell times, overall representing the rate of protein synthesis (Fig. 8o). Loss of *Ire1α* leads to a decreased number of translation sites (Fig. 4k) and slower translocating ribosomes (Fig. 4h), which is associated with a global reduction of HPG incorporation, indicative of lower rate of protein synthesis.

Seemingly, on top of elegant transcriptional regulation (*14, 15*), generation of upper layer neurons relies on specific pathways for translational control. According to our data, specific features of 5’UTR regions in genes critical for upper layers (*50*) require activity of translation initiation complexes, comprising eIF4A1, which unwind cap-proximal regions of mRNA, especially ones with G4s, prior to its ribosomal loading (*51*). In line with this are our observations on decreased translation rates upon *eIF4A1* KO (Fig. 4c and 4d) associated with defects in upper layer neurogenesis (Fig. 3g and 3h). Loss of *eIF4A1* or *Ire1α* leads to a decreased translation efficiency of Satb2 reporter. On the other had, CTIP2 reporter in these cases is translated more efficiently, indicative of ribosome preference redistribution, dependent on 5’UTR of fate determinants (Fig. 8 and S6).

The *Ire1α* cKO partially mimics the *Satb2* KO (*8*), with the loss of upper layer type neurons at the expense of CTIP2-expressing deeper layer cells (Fig. 1 and S2). Strikingly, in the *Ire1α* cKO cortex CTIP2-expressing neurons are specified during late neurogenesis (Fig. S2e-S2h). Wild type neuronal progenitors at e12.5 generate CTIP2 and Satb2 positive cells, and as development progresses, late progenitors give rise to Satb2-expressing progeny (Fig. S2 and 5-6). Such progressive restriction of neuronal progenitor potency is abolished upon *Ire1α* loss-of-function. In contrast, high activity of IRE1α correlates with lower expression of stemness markers, like SOX2, POU3F2 or OLIG2 (*52*).

It is worth emphasizing the results in Fig. 5 and 6 represent different ways of lineage labeling (bulk nucleofection versus clonal progenitor tree), which accounts for different fate signatures in the postmitotic progeny. Our approach to mix labeled cells derived from embryonic cortices of different developmental stages allows for exposure to identical culturing conditions. An interesting future direction is to examine how cellular and environmental composition affects the identity of progenitors, neurons, and associated translation rates.

Global attenuation of translation during the UPR is linked to PERK-mediated phosphorylation of eIF2α at Ser51 (*53*). The stress response alters the cellular translation machinery and its affinity towards open reading frames of specific type, position, and secondary structures (*54, 55*). We report that loss of *Ire1α* results in lower rates of protein synthesis in neuronal progenitors and neurons, independently of eIF2α, or its canonical downstream splicing client Xbp1 (Fig. S4a and S4b). *Ire1α* regulates the amount of translating polysomes, with their higher abundance in the cKO. In the context of lower translation flux, this represents ribosome stalling and/or slower elongation (Fig. 4h), reminiscent of observations in *Huntingtin* mutants (*40*). As Ire1α binds 80S ribosomes directly with high affinity [Fig. 3j and (*25*)], and is in the vicinity of the ribosomes embedded in the ER by the translocon, it is well positioned to impact protein synthesis directly during cortex development. The interaction between Ire1α with small ribosomal subunit is strongest at E12.5 (Fig. 3j), when multipotent progenitors able to generate upper and deeper neuronal lineages are present in the cortex.

Protein synthesis inhibition with CHX resembles the *Ire1α* cKO and *eIF4A1* KO (albeit engendering expected stronger phenotypes, given the global inhibition of translation by the compound), with defects in Satb2 expression and axonal targeting of Tau-1, leading to the formation of symmetrical, non-polarized neurons and ablation of Satb2 (Fig. 7 and S7). Both the loss of *Ire1α* and CHX spare the expression and subcellular localization of deep layer neurons-associated CTIP2 and dendrite-enriched MAP2, indicative of the specific requirement for Ire1α and high translation rates during upper layer neurogenesis and polarization. In line with this, we find that late neuronal progenitors display a drastic increase in translation rates, which represent the required molecular background for efficient eIF4A1-dependent unwinding of 5’UTR of Satb2. When protein synthesis rates are lower, ribosomes promote translation of deeper layer determinants.

Apart from being a site of protein synthesis, the ER has been also reported as an organelle crucial for neuronal polarization (*56*). Upon *Ire1α* KO, we report a mislocalization of the ER to the polarizing neurite (Fig. 2m and 2n). Defects in axon specification, and thus axonal targeting of Tau-1, AnkG and Na_v_s (Fig. S3) might indicate ER dysfunction upon *Ire1α* loss, implicating its role in ER integrity and neuronal polarization (*57*). Loss of polarization might also partially explain the filamin A-dependent *Ire1α* loss-of-function phenotype in laminar positioning of cortical neurons (*23*). Emergence of either multiple axons or more commonly no axons in *Ire1α* KO neurons (Fig. 2) implies a disruption of distinct axon polarity-regulating processes in this mutant, likely involving cellular sorting of axonal proteins and ER-dependent microtubule stability, overall leading to disturbed transmission of action potentials. Cortical neuronal migration and neuronal polarization involves *Ire1α-*driven control of multiple downstream processes, filamin A anchoring on one hand, eEF-2- and eIF4A1-regulated translation on the other hand.

Taken together, our study reinforces the powerful impact of post-transcriptional mechanisms in cortex development. Translational regulation of gene expression during neuronal specification is layered on top of transcriptional mechanisms, many of which have been highlighted in excellent recent work (*14, 15*). Modulation of translation rates may be a particular requirement for upper layer specification and morphology, evolving as a mechanism to specify upper layers in the late stages of development while restricting the potential of neural progenitors to this lineage. We speculate that a swift upregulation of translation flux in late progenitors might be necessary to efficiently synthesize determinants of upper layer neurons. In the light of eIF4A1 requirement for Satb2 identity, we demonstrate that 5’UTR-embedded structural features are present in mRNAs critical for differentiation of upper layer neurons. Given developmental downregulation of Ire1α, it is plausible that other translational regulators alongside Ire1α drive the increased translation efficiency in upper layer progenitors. Thus, translational control by in part by Ire1α is deployed in normal development to promote upper layer neurogenesis, generate their axons, and wire the mammalian cortex.

## SOURCE DATA

## ACKNOWLEDGEMENTS

This work was supported by the German Research Foundation (TA 303/15-1, to VT) and the Russian Scientific Foundation (21-65-00017, 19-34-51009, to VT). We thank Mengfei Gao, Helge Ewers, the organizers of CAJAL Advanced Imaging Methods Course for Cellular Neuroscience and Boehringer Ingelheim Fonds for the introduction to expansion microscopy. We thank Paraskevi Bessa-Newman and Ekaterina Epifanova for the useful tips, Hiroshi Kawabe for the pRai plasmid, Frederick Rehfeld for dsRed expression vector and the Charité Viral Core Facility for production of viral particles, and Nils Brose for his continuous support and advice.

## AUTHOR CONTRIBUTIONS

M.C.A. carried out and conceptualized the study, with significant contribution from E.B. V.T. designed the inhibitor screening, initiated, and supervised the project. A. G. N. performed the experiments with RNAseq and the bioinformatics. M. L. K. purified the ribosomes ex-vivo and analyzed their profiles. T. S. performed the molecular cloning. R. D. significantly contributed to histological and cell culture experiments. M. B. recorded from autaptic cultures and analyzed the results. P. T. shared his expertise and protocols for the FUNCAT and click chemistry. D. K. helped with the initial screening and share the molecule library. O. J. performed proteomic analyses. T. I. shared the *Ire1α*^f/f^ mouse line. M. R., C. M. T. S., and C. R. supervised works in their labs.

## CONTACT FOR REAGENTS AND RESOURCE SHARING

Further information and requests for resources and reagents should be directed to Victor Tarabykin. Detailed information on the materials used in this study can be found in Table S3.

## EXPERIMENTAL MATERIAL AND SUBJECT DETAILS

### Animals

Mouse (*Mus musculus*) lines described in this study have been maintained in the animal facilities of the the Charité University Hospital and Lobachevsky State University. Wild type mice were of NMRI strain. In *Ire1α*^f/f^ mouse line (*31*), exons 20-21 of *Ire1α* were engineered to be flanked by loxP sites. To inactivate *Ire1α* in the developing cortex, we crossed *Ire1α*^f/f^ mice with the *Emx1*^Cre/+^ line, in which Cre recombinase is expressed from the *Emx1* gene allele (Gorski et al., 2002). As described in our previous work, for the breeding we exclusively used *Ire1α*^f/f^ males and *Ire1α*^f/f^; *Emx1*^Cre/+^ females to circumvent leaky expression of Cre (*58*). Animals homozygous for the loxP alleles in both lines were viable, fertile, and born at the expected Mendelian ratio, and exhibited no overt phenotypic changes in the cage environment. For experiments with tdTomato reporter (Fig. 1a-1b), *Satb2*^Cre/+^ (*27*) males were mated to NMRI wild type females. The date of vaginal plug was counted as E0.5. All mice were sacrificed by administering a lethal dose of pentobarbital.

All experiments were performed in compliance with the guidelines for the welfare of experimental animals approved by the State Office for Health and Social Affairs, Council in Berlin, Landesamt für Gesundheit und Soziales (LaGeSo), permissions G0079/11, G206/16, and G54/19, and by the Ethical Committee of the Lobachevsky State University of Nizhny Novgorod.

### Sex and age/developmental stage of animals for *in vivo* experiments

Littermates of both sexes were randomly assigned to experimental groups during experimental procedures or collection of embryonic tissue. Developmental stages or stages at experimental interventions are listed on the figures or in the figure legends.

### Murine primary neurons

Embryonic brains were dissected on ice-cold Hank’s Balanced Salt Solution (HBSS) under stereomicroscope to collect cortices in 5 mL of ice-cold 0.25% trypsin (Gibco) solution. After incubation for 20 min at 37°C with occasional swirling, digestion was terminated by addition of 1 mL of fetal bovine serum (FBS, Gibco) and 1U of DNaseI (Roche) was added to the tube for 1 min. Next, cortices were carefully washed twice with 5 mL of pre-warmed (37°C) Complete Neurobasal medium. Further, cortices were carefully triturated in 1 mL Complete Neurobasal medium 10 times using P200 pipette tip. The debris were then let sink for 1 minute and 150 μL the supernatant transferred to 900 μL of fresh Complete Neurobasal at 37°C. After second round of trituration, cells were counted with Naubauer counting chamber and neurons were seeded in 24 well-plate formats at 120 000 cells (EUE) or 240 000 cells (after nucleofection) per coverslip in 1mL of Complete Neurobasal medium. The day of neuronal prep was counted as day *in vitro* 0 (DIV0). Neurons were cultivated at 37°C in the presence of 5% carbon dioxide in HERA-cell240 (Heraeus) incubator. For the experiments with DMSO and CHX, neuronal cultures were treated with the substance two hours post-plating. Given substance was thoroughly mixed with fresh medium and placed on attached cells.

Autaptic hippocampal cultures were prepared from P0 *Ire1α*^f/f^ mice as described (*59*).

<u>Complete Neurobasal</u>: 500 mL Neurobasal (Gibco, Life Technologies), 10 mL B-27 (Gibco, Life Technologies), 5 mL GlutaMAX (Gibco, Life Technologies), 5 mL penicillin/streptomycin (Life Technologies, Gibco), 1 mL Primocin (Invitrogen).

### Mouse embryonic fibroblasts (MEFs) from *Ire1α*^f/f^ mice

MEFs were collected from E13.5 *Ire1α*^f/f^ embryos. Briefly, embryo body was dissected, minced with razor blade and incubated with 1mL 0.25% trypsin (Gibco) at 37°C for 30 minutes. Trypsinization reaction was terminated by addition of 4 mL MEF medium and the tissue was triturated 20 times to release single cells. Cellular suspension was transferred to T75 flask with 15 mL of fresh MEF medium and cells were incubated until confluency. Cells were split according to standard cell culture protocols. Cell cultures were expanded and frozen at passage five, flash frozen in liquid nitrogen and stored until use at −80°C. Before each experiment, correct genotype of the fibroblasts was validated using the DNA isolated from the cell suspension according to the genotyping protocol for *Ire1α.* MEFs were infected using AAVs according to standard infection protocols.

<u>MEF medium</u>: 500 mL DMEM (Life Technologies), 50 mL FCS (Gibco), 5 mL penicillin/streptomycin (Life Technologies, Gibco), 5 mL GlutaMAX (Gibco).

## METHODS

### Lentivirus production and infection of autaptic hippocampal cells

Production of lentiviral particles and infection of autaptic cultures was performed by the Charité Viral Core Facility according to standard published protocols.

### Electrophysiological Recordings

Voltage and current clamp experiments were performed as described before (*59*).

### Expression vectors

For our screening, we used beta-actin driven expression constructs pCAG-EGFP and pCAG-flox-stop-flox-tdTomato, which are described in our previous work in detail (*26*)

### Cloning strategies for constructs generated in this study

Cloning of all expression vectors in this study was performed using the NEBuilder system according to manufacturer’s protocol (New England BioLabs). DNA fragments were amplified using GXL Prime Star DNA polymerase (Takara) using cDNA libraries as templates. Destination vectors were linearized with EcoRI-HF (New England BioLabs).

#### pRai-HA-IRE1α

Human IRE1α cDNA (NM_001433.3) was amplified from plasmid template [Addgene, #13009, (*60*)] using the following oligos: 5’-agattacgctatctgtacaggcATGCCGGCCCGGCGGCTG-3’ and 5’-ggccgctagcccgggtaccgCTTGGTTTGGGAAGCCTGGTCTCCCTGC-3’ and inserted into the modified pRaichu vector (*61*).

#### pCAGIG-6XHis-eEF2

Murine eEF2 cDNA (NM_007907.2) was amplified from cDNA library using the following oligos: 5’-gtctcatcattttggcaaagATGCATCATCATCATCATCATGTGAACTTCACAGTAGATC-3’ and 5’-cggccgcgatatcctcgaggCTACAGTTTGTCCAGGAAGTTG-3’ and inserted into pCAGIG vector for simultaneous expression of 6XHis-tagged eEF2 and EGFP.

#### pCAG-5’UTR-Satb2 and -CTIP2 reporter

Satb2 and CTIP2 5’UTR sequences were amplified using a mouse embryonic cDNA library and Prime Star GXL polymerase. An existing pCAG-IRES-tdTom vector was linearized using EcoRI digestion and PCR-amplified fragments were inserted and fused together with a GFP coding sequence. Primers for PCR had the following sequences: 5’UTR Satb2_fwd, 5’-gtctcatcattttggcaaagCGCCCCCATCATCATAAC-3’; 5’UTR Satb2_rev, 5’-ccttgctcacCATGCTGCTCCGATTTGG-3’; GFP_fwd, 5’-gagcagcatgGTGAGCAAGGGCGAGGAG-3’; GFP_rev, 5’-cggccgcgatatcctcgaggTTACTTGTACAGCTCGTCCATG-3’; 5’UTR CTIP2_fwd, 5’-gtctcatcattttggcaaagAATTTATTTTAGCCTTTTCTCTATTTTAGAGCAAG-3’; 5’UTR CTIP2_rev, 5’-ccttgctcacCATTGCCCCGGCATCTATTC-3’; GFP_fwd, 5’-cggggcaatgGTGAGCAAGGGCGAGGAG-3’; GFP_rev, 5’-cggccgcgatatcctcgaggTTACTTGTACAGCTCGTCCATG-3’. The fragments were recombined with the vector backbone using NEBuilder system. After that, the 5’UTR fragments fused to GFP and IRES sequence were shuttled to a pCAG vector (*62*) and fused with dsRed encoding sequence also using NEBuilder system.

### Nucleofection of primary neurons

Transfection of primary cortical cells was performed using nucleofection (Lonza Bioscience) according to our previously published protocol (*26*).

### Small molecule inhibitor screening

Right after trituration, E13.5 embryonic cortical neurons prepared from *Satb2*^Cre/+^ mice, were nucleofected with pCAG-EGFP and pCAG-loxP-Stop-loxP-tdTomato plasmids and seeded at 120 000 cells per well of 96-well plate. Two hours post-plating, cultures were treated with compounds at two concentrations, in technical duplicates. Cells were then cultivated until DIV2, when the proportion of *Satb2*^tdTom^ neurons normalized to EGFP positive cells was determined using FACS as described (*26*).

### *Ex Utero* Electroporation (EUE)

Embryonic brains were injected with a glass capillary to fill the entire volume of a single lateral embryonic ventricle with DNA (final concentration of 500 ng/μL), followed by electroporation using 6 pulses of 35V applied using platinum electrodes. The isolated heads were then placed in ice cold HBSS until proceeding with primary cultures.

### Immunocytochemistry

Cells cultured on coverslips were fixed with cold (4°C) 4% PFA, 4% sucrose (Merck) in PBS for 20 minutes in room temperature, washed three times with PBS and incubated with Blocking buffer for 30 minutes. Next, cells were incubated with primary antibodies diluted with blocking buffer accordingly for 16-20 hours at 4°C with moderate shaking, followed by washing 3 times with PBS. Further, secondary antibodies coupled to appropriate fluorophore diluted in blocking buffer were applied for 2 hours at room temperature. After washing 3 times with PBS, coverslips were briefly rinsed with ddH_2_O and mounted on Superfrost Plus glass slides (Thermo Scientific) with Immu-Mount mounting medium (Shandon, Thermo-Scientific).

<u>Blocking buffer</u>: 5% horse serum (Gibco, Life Technologies), 0.3% Triton X-100 (Roche), in PBS buffer

### Fluorescent Noncanonical Amino Acid Tagging (FUNCAT)

For the fluorescent labeling of newly synthesized proteins, we modified previously published methods (*63*). In brief, neuronal cultures were fed with 1mM HPG-supplemented Methionine-free Neurobasal Medium, at DIV1 and DIV5, for the time indicated on the figures and in the figure legends. Cells were then extensively washed in PBS prior to fixation with 4%PFA/4% sucrose for 20 min. Control and HPG fed neurons were then fluorescently labelled with 1µM Sulfo-Cyanine5 azide (Lumiprobe). To facilitate azide-alkyne binding we applied sCy5az in a PBS based “click solution” containing 0.2 mM Tris(3-hydroxypropyltriazolylmethyl) amine (THPTA), 20 mM sodium L-ascorbate, and 0.2 mM copper (II) sulfate pentahydrate (all from Sigma-Aldrich). Cells were incubated at room temperature for 20 minutes, before PBS washing and immunocytochemistry (described above).

<u>Met-free Neurobasal Medium</u>: 500 mL Met-, Lys-, Arg-free Neurobasal (Gibco, Life Technologies), 10 mL B-27 (Gibco, Life Technologies), 5 mL GlutaMAX (Gibco, Life Technologies), supplemented with final 1 mM L-HPG (Jena), 0.8 mM Lys, and 0.4 mM Arg.

### Puromycin metabolic labeling

To metabolically label proteins using puromycin, we used DIV5 *Ire1α*^f/f^ mouse embryonic fibroblasts from infected with AAV particles (Charité Viral Core Facility) encoding for EGFP (control) and EGFP-p2a-Cre at DIV0. Cells were incubated with 20 μg/mL puromycin for 10 minutes, as described (*45*). The cells were washed with PBS and lysed in 1X Laemmli sample at 70°C for 20 minutes. Proteins were then resolved in SDS-PAGE and analyzed using Western blotting with anti-puromycin antibody. The puromycin incorporation was normalized to the protein loading amount.

### Ribosome run-off assay

To quantify ribosome run-off, we used DIV5 *Ire1α*^f/f^ mouse embryonic fibroblasts from infected with AAV particles (Charité Viral Core Facility) encoding for EGFP (control) and EGFP-p2a-Cre at DIV0. Cells were incubated with 2 μg/mL harringtonine for indicated time and immediately after with 20 μg/mL for 10 minutes. Next, cells were washed with PBS and analyzed as described for “Puromycin metabolic labeling”. Puromycin incorporation was normalized to the protein loading amount and to the incorporation in Control at 1 min.

### SunTag

To study translation dynamics using the SunTag system, we used DIV5 mouse embryonic fibroblasts from *Ire1α*^f/f^ infected with AAV particles (Charité Viral Core Facility): empty virus (control) and one encoding for Cre. At DIV4, MEFs were transfected with SunTag labeling plasmids including pcDNA4TO-24xGCN4_v4-BFP-24xPP7 (Addgene 74929), pHR-scFv-GCN4-sfGFP-GB1-NLS (Addgene 60906), and pHR-tdPP7-3xmCherry (Addgene 74926). After 20 hours, the reporter expression was induced with 1 μg/mL doxycycline (Sigma) for 1 hour (*46*). Next, cells were fixed, mounted on SuperFrost slides and fluorescence was detected using spinning disc microscopy.

### Isolation of RNA, cDNA library preparation and RNA sequencing

Tools and work surfaces used for RNA work were thoroughly cleaned with 70% ethanol (Sigma) and rinsed with RNA-Zap (Thermo Fisher) before the procedure. Embryonic brains at E18.5 were dissected in DEPC-PBS and flash-frozen in liquid nitrogen until the purification procedure. RNA was isolated using RNeasy columns (RNeasy Mini Kit, Qiagen). Further steps were performed according to the manufacturer’s protocols. RNA was eluted from the silica membranes using 30 μL of molecular biology-grade (MB)-H_2_O. Quality and concentration of the prepared RNA was determined using an Agilent Bioanalyzer. Single or paired end TruSeq Stranded total RNA libraries were made with Ribo-Zero Gold rRNA depletion and sequenced using Illumina Nextseq 500/550. Raw reads were aligned to GRCm38.p5 using STAR 2.5.3a (*64*) with parameters --outFilterMultimapNmax 100 -- winAnchormultimapNmax 100 --outSamstrandField intronMotif and the gencode vM16 basic annotation transcriptome. A single count table was generated from the resulting bam files using TETranscripts (*65*). Alignment pipeline was executed using Snakemake (https://snakemake.github.io/). Counts were then fit to a quasi-likelihood negative binomial generalized log-linear model and tested for differential expression using EdgeR (*66*).

### Biochemical experiments

Details of the protocols used for Preparation of cortical lysates, Protein concentration measurements, Sodium dodecyl sulfate polyacrylamide gel electrophoresis (SDS-PAGE), Western blotting, Quantification of protein levels are described in our previous works (*18, 61*).

### Co-Immunoprecipitation

Mouse cortex tissues were homogenized in E1A lysis buffer (150 mM NaCl, 50 mM HEPES, pH 7.0), supplemented with protease inhibitor cocktail (Roche), 2.5 mM sodium pyrophosphate, 1 mM beta-glycerophosphate, 10 mM sodium fluoride, 1 mM Na_3_VO_4_, and 1 mM PMSF (Millipore). For a single assay point, four cortices of E12.5 and two of E14.5 were lysed together. The lysate was centrifuged at 12 000 g for 15 min and the supernatant with 0.25 mg total protein was incubated with 0.5 μg anti-Ire1α (Cell Signaling) antibody for 4 hours at 4°C. Next, the supernatant was incubated with 25 μL Sepharose A beads (Roche) for 1 hour at 4°C. As a control, the supernatant was incubated with 0.5 μg rabbit IgG (Cell Signaling). After extensive washing (three times, buffer E1A), the bound proteins were eluted in 1X Laemmli sample buffer and incubated at 70°C for 20 minutes, prior to SDS-PAGE and Western blotting. To quantify the strength of interaction between Ire1a at E12.5 and E14.5, the density of indicated proteins were normalized to the density of Co-IPed Ire1α and to the strength of interaction at E12.5.

### Polysome purification

Ribosome fractionation and polysome purification from E18.5 cortex was performed as published protocol (*36*). Seven brains (14 neocortex hemispheres) of each genotype were pooled per biological replicate, with analysis in two replicates per condition.

### Cryosectioning

For all histological procedures, brain sections were prepared on Leica CM3050S cryostat. Prior to cryosectioning, brains were passed through a sucrose gradient to cryoprotect the tissue. Next, brains were frozen in −38 to −40°C isopentane (Roth). Coronal cryosections of 50 μm thickness were collected in PBS/0.01% sodium azide solution. For *in situ* hybridization 16 μm thickness sections were collected.

### Fluorescence *in situ* hybridization (FISH)

We used RNAscope Technology to detect mRNA of Ire1α according to the manufacturer’s protocols (ACD). Prior to hybridization, embryonic brains at E12.5, E14.5 and E18.5 were collected in DEPC-PBS and incubated in 4%PFA/PBS/DEPC for 16-20 hours at 4°C. Brains were then passed through a series of sucrose solutions (10%-20%-30%/PBS) until they reach osmotic equilibrium, embedded in O. C. T. Compound (Tissue-Tek) in a plastic cryoblock mold and frozen on dry ice. Coronal sections of 16 μm thickness were collected using the cryostat.

### Immunohistochemistry

Fixed brain sections were washed with PBS three times at room temperature prior to the procedure to remove the sucrose and freezing compound residue. The sections were then incubated with Blocking solution for one hour at room temperature, then with the primary antibody and DAPI diluted in blocking buffer for 16-20 hours at 4°C, washed in PBS three times for 30 minutes and incubated with secondary antibody diluted in the blocking buffer for up to four hours at room temperature. Next, sections were incubated with PBS for 30 minutes three times and mounted with cover glass (Menzel-Gläser) and Immu-Mount mounting medium (Shandon, Thermo-Scientific).

<u>Blocking solution</u>: 5% horse serum, 0.5% (v/v) Triton X-100, PBS

### BrdU Injection and Immunohistochemistry

Pregnant females at appropriate stages were intraperitoneally injected with BrdU, as described (*6*). Brains were then isolated at p2 from mice of both genotypes (Control: *Ire1α*^f/f^ and cKO: *Ire1α*^f/f^; *Emx1*^Cre/+^), fixed in 4% PFA, passed through sucrose gradient and frozen in isopentane bath. Free floating cryosections of 50 μm thickness were placed on the SuperFrost Plus glass slides and let dry overnight at +4°C. Antigen retrieval was performed in boiling citrate-based Antigen Unmasking Solution (Vector, pH 6.0) for seven minutes in a microwave followed by 20 min of cooling on ice. Next, sections on slides were immunolabeled according to the protocol.

### Expansion Microscopy

For the purpose of expansion of coronal brain sections to visualize morphology of EGFP-expressing neurons, we followed previously published protocol (*67*). For immunohistochemistry, we used anti-EGFP antibody (Rockland) at 1:200 and the secondary Alexa488-coupled one at 1:100. After PBS washes, the sections were incubated with AcX crosslinker (10 mg/mL) in 150 mM NaHCO_3_ for 20 hours at room temperature, washed with PBS, incubated for one hour with monomer solution (19% sodium acrylate, 10% acrylamide, 0.1% bis-N’,N’-methylene-bisacrylamide, 0.01% 4-hydroxy-TEMPO in PBS) and gel was polymerized by addition of 0.2% APS/0.2% TEMED for two hours at 37°C. The section embedded in the gel was then incubated with a proteinase K in a digestion buffer (50 mM Tris-Cl, 800 mM guanidine-Cl, 2 mM CaCl_2_, 0.5% Triton X-100 in pH 8.0) for 20 hours at 37°C with gentle agitation. Next, section was transferred to a Petri dish with ddH_2_O, which was changed at least five times for two days. The expansion factor was determined before mounting of the specimen. Sections were then transferred onto the glass-bottom chambers (Ibidi) and imaged using Leica Sp8 confocal.

### *In utero* electroporation (IUE)

Prior to IUE, DNA was diluted in TE buffer and mixed with 0.1% Fast Green FCF (Sigma-Aldrich). Final concentration of DNA used for transfecting cortical progenitors was 200-500 ng/μL. The details on the procedure are exactly as we described before (*18*).

### Antibodies

For Western blotting, all antibodies were diluted 1:750, apart from anti-β-actin (used at 1:2000), and anti-puromycin (used at 1:10000), in 5% non-fat dry milk in TBS-T buffer (TBS buffer with 0.1% Tween-20). Antibodies against phospho-proteins, anti-puromycin and anti-Ire1α were diluted in 3%BSA in TBS-T buffer. Secondary antibodies coupled with HRP were diluted 1:10000 in the same buffer as the primary ones.

For immunocytochemistry, all primary antibodies were diluted 1:300 in the blocking buffer, and the appropriate fluorophore-conjugated secondary antibodies were used at 1:750.

For immunohistochemistry, primary antibodies and the respective fluorophore-linked secondary ones were diluted 1:500 in the blocking buffer.

### RNA structure prediction and quantification

The sequences of murine 5’UTR of Satb2 and CTIP2 mRNA were identified using the UCSC Genome Browser (Mouse Assembly GRCm39/mm39). The structures of 5’UTRs were modeled using the RNA Web Suite http://rna.tbi.univie.ac.at/cgi-bin/RNAWebSuite/RNAfold.cgi. The G-quadruplexes and R-loops were detected and quantified using the G4Hunter web application, https://bioinformatics.ibp.cz/#/ (*68*).

### Data analysis

Details concerning Quantitative Western blotting, Axon counting assays *in vitro*, Neuronal morphometry, Quantification of distribution of cortical neurons and polarity classification are described in detailed in our previous publications (*18, 61*).

### Confocal imaging of immunostaining signals and analysis

Images of primary cell cultures and brain sections after immunostaining were acquired using confocal Leica SL, Leica Sp8 or Zeiss Spinning Disc Microscope.

### Colocalization with identity markers

Neuronal identity on brain sections was quantified as fraction of all EGFP-expressing neurons. Identity of primary cells on coverslips was determined as fraction of fluorescent reporter-expressing cell. Based on the numbers of dsRed-or EGFP-expressing cells, we either quantified at least 4 fields of view for high-density cultures, or imaged a given number of cells (indicated on the bar graphs or Table S1) and determined the number of cells expressing a marker protein for cultures with sparse labeling, i.e. after EUE. Compare Fig. S7.

### Laminar positioning (% of CP)

We based our positioning analysis on previous published reports (*16*). To determine the position of neurons in the cortical plate, confocal images of EGFP signals were first transformed so that the pia is perpendicularly oriented to the horizontal axis. Next, positions of neurons were marked in Fiji using the Cell Counter plug-in. Using the y-coordinate, we then expressed the position of a given cell relative to the size of the CP and plotted it as % CP, with 0% being the bottom of the cortical plate / the subplate (SP) and a 100% being the pial surface / the marginal zone (MZ). Positions of all cells were then plotted individually on the graph. Only brains with comparable electroporation efficiencies were analyzed.

### Quantification of fluorescence intensity

For quantifications of HPG-647 fluorescent intensity, we defined a region of interest based on the fluorescent marker, i.e. dsRed or EGFP signals. Using Fiji software, the raw integrated density was then normalized to the cell surface area of the outlined cell compartment. For our quantifications, we imaged the primary cell cultures as Z-stacks and for the analysis, we used maximum intensity projections. For quantification of EGFP fluorescence intensity in cells electroporated with 5’UTR reporter, integrated intensity was quantified, given homogenous cell size. For the experiments in fibroblasts transfected with the reporter constructs, because of rather variable cell sizes, mean fluorescence in a defined cell area was plotted. For the SunTag analysis, mean EGFP fluorescence intensity in a given mCherry puncta was quantified using the Cell Counter plugin.

### Gene Set Enrichment Analysis (GSEA)

GSEA was performed on total counts from cKO and control polysome fraction RNAseq in classic mode, testing set, the difference of classes metric, over 1000 permutations with the c5.go.mf.v7.2.symbols.gmt (molecular function) annotation. This gave 406/772 gene sets upregulated in cKO polysomes where 158 gene sets were significant at FDR<25%. Detailed results are presented in the Table S2.

### Statistical analyses

All statistics were performed using Prism Graph Pad software. Detailed information on statistics can be found in the supplementary material. Description of statistical tests, definition of center, dispersion, precision and definition of significance are also listed in the figure legends. Briefly, distribution of datapoints was determined using D’Agostino and Pearson normality test. For normally distributed data, we used unpaired two-tailed t-test, or one/two-way ANOVA with Bonferroni post hoc test; for data with not normal distribution, we used Mann-Whitney test to compare two groups, or Kruskal-Wallis test with Dunn’s Multiple comparison test for comparison between multiple groups, unless specified otherwise. For detailed information on all numerical values, sample size and statistical tests used in all experiments, see Table S1.

## SUPPLEMENTARY TABLES

Table S1. Numerical values, sample size and statistics for experiments in this paper.

Table S2. GSEA report for RNA upregulated in cKO polysomes.

Table S3. Key resources used in this work.

---

**Figure S1.**
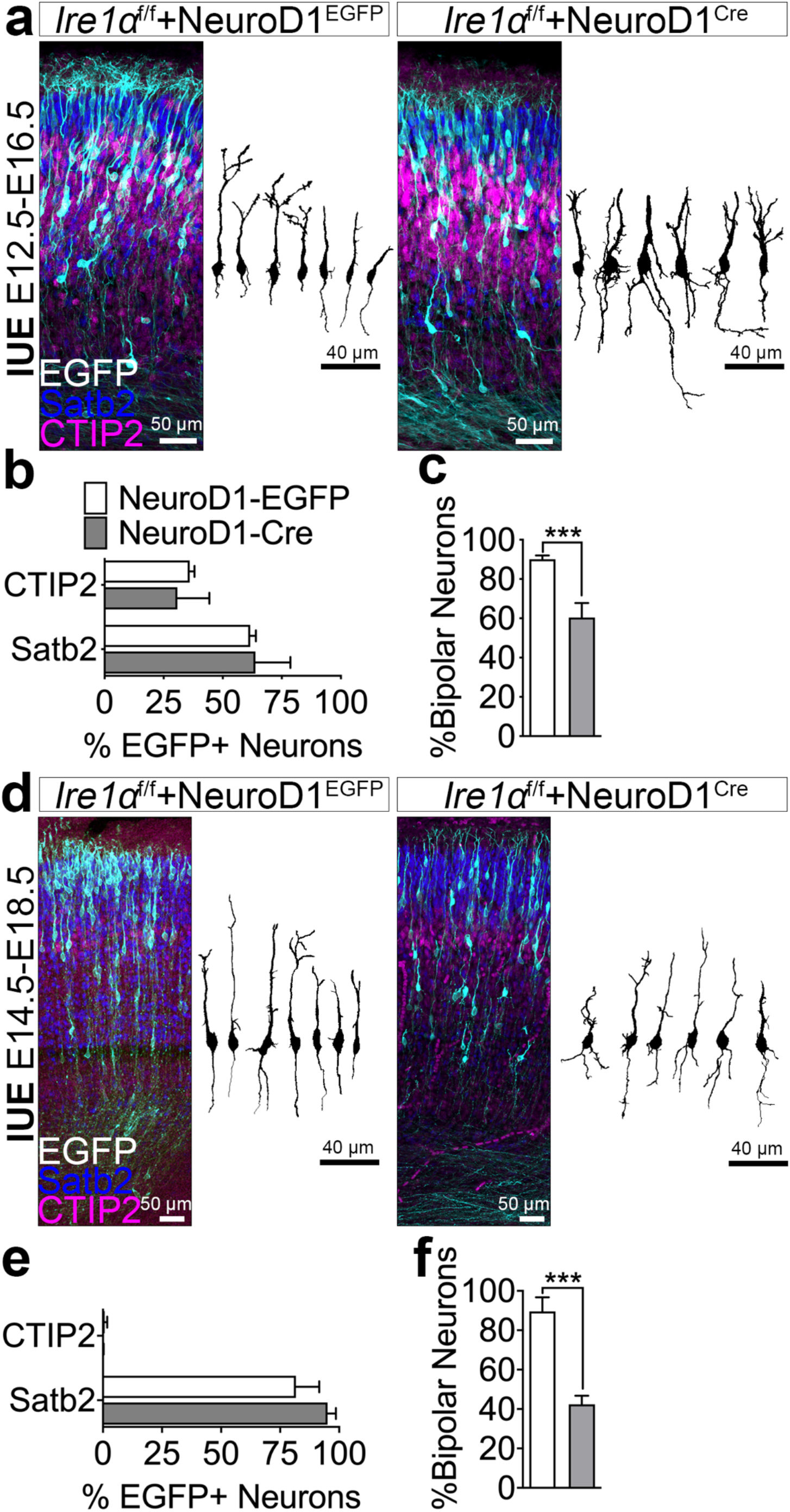
Related to Fig. 1. Ire1α-driven acquisition of the upper layer identity, but not of bipolar morphology, is embedded in neuronal progenitors. (a, d) Representative images of immunostaining in coronal cortical sections from E16.5 (a) or E18.5 (C) *Ire1α*^f/f^ embryos after IUE at E12.5 (a) or E14.5 (d) with plasmids encoding for EGFP under the promoter of *NeuroD1* gene or Cre under the promoter of *NeuroD1* together with loxP-Stop-loxP-EGFP. Shown are also representative, EGFP-based tracings of single neurons. (b, e) Quantification of neuronal cell identity in (a) and (d). (c, f) Average proportion of bipolar neurons in (a) and (d). Bar graphs show averages ± S.D. For statistical analyses, D’Agostino-Pearson normality test and Mann-Whitney test. *** p < 0.001.

**Figure S2.**
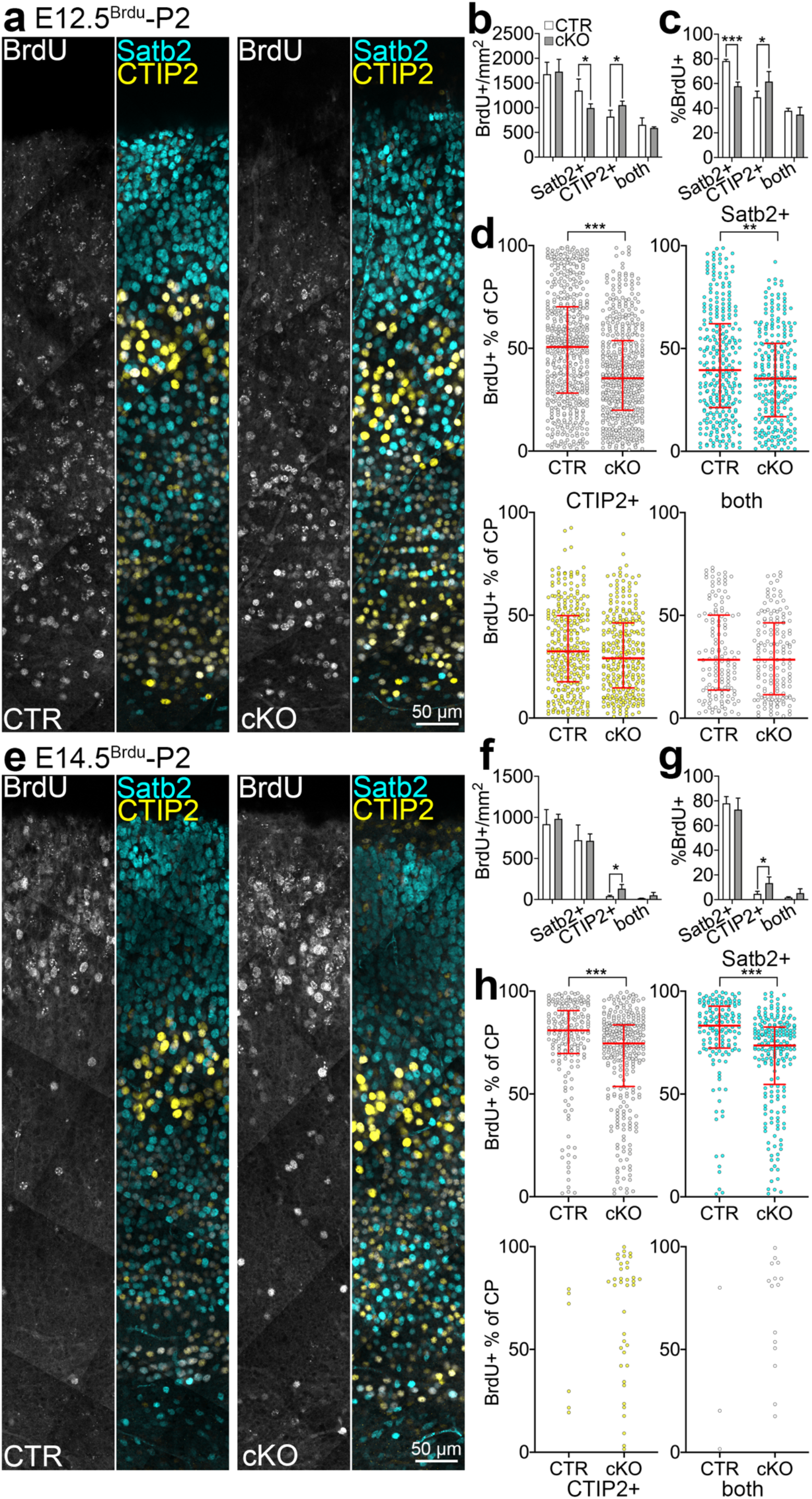
Related to Fig. 1. Birth dates and laminar positions of Satb2+ and CTIP2+ neurons in *Ire1α* cKO cortex. (a, e) Representative images of immunostaining in control or cKO coronal cortical sections after BrdU injection at E12.5 (a-d) or E14.5 (e-h). (b), (f) Quantification of cell density expressing an indicated nuclear stain. “Both”, cells co-expressing of Satb2 and CTIP2. (c), (g) Quantification of the proportion of neurons expressing indicated marker within all BrdU+ cells. (d), (h) Laminar distribution of neurons expressing indicated marker. The position of each neuron was normalized to the thickness of the CP and represented as a dot (100% CP – pia, 0% CP – bottom of the CP). Graph contains pooled data from indicated number of brains (Table S1). Bar graphs show averages ± S.D. Line and error bars on (d) and (h) indicate median and interquartile range. For statistical analyses, D’Agostino-Pearson normality test and (d) and (h), Mann-Whitney test; (b-c) and (f-g) unpaired t-test. 0.001 < ** p < 0.01; *** p < 0.001.

**Fig. S3.**
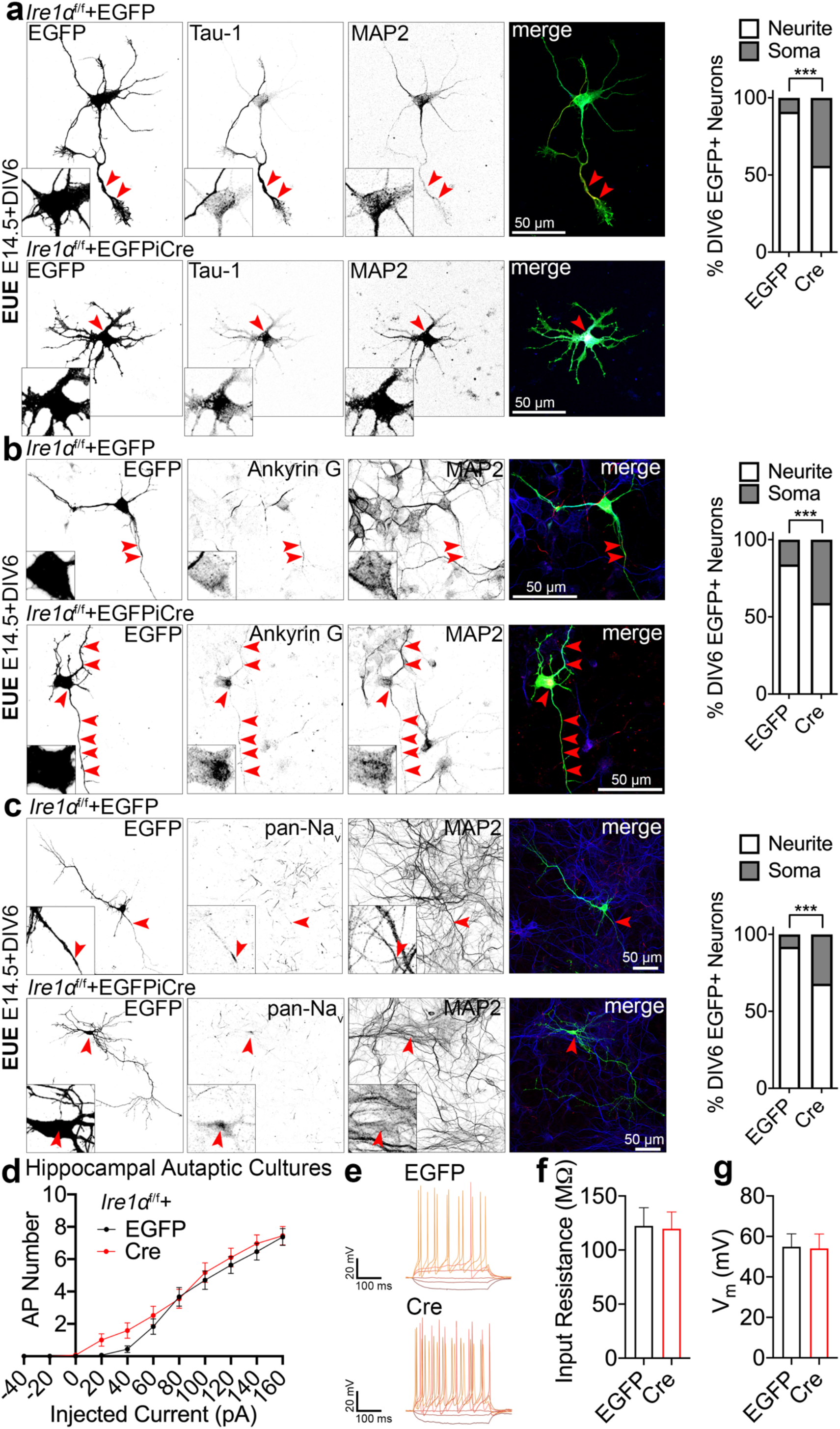
Related to Fig. 2. Loss of *Ire1α* disrupts axon initial segment and current responses. (a-c) Images of representative DIV6 primary cortical *Ire1α*^f/f^ neurons after EUE at E14.5 for EGFP or EGFP and Cre expression. Neuronal cultures were immunolabeled for indicated axonal and dendritic marker proteins. Graphs show quantification of the subcellular distribution of axonal markers. *Ire1α-*deficient neurons display somatic accumulations of Tau-1, Ankyrin G and Na_v_. Red arrowheads indicate enrichment of axonal markers. Insets on the bottom left of black and white images are zoom-ins to the somata or places of axonal marker accumulation. (d-g) Electrophysiological recordings from autaptic hippocampal cultures prepared from P0 *Ire1α*^f/f^ pups. Neuronal cultures were infected with lentiviruses encoding for EGFP or Cre. (d) Quantification of the number of APs generated after injection of increasing amount of current. (e) Example traces of recordings described in (d). Average input resistance (f) and holding potential (g). Data points and error bars on (d) represent mean ± S.E.M. and graphs on (f-g), mean ± S.D. For statistical analyses, (a-c), Fisher’s test; (d), one-way ANOVA; (f-g), Mann-Whitney test. *** p < 0.001.

**Fig. S4.**
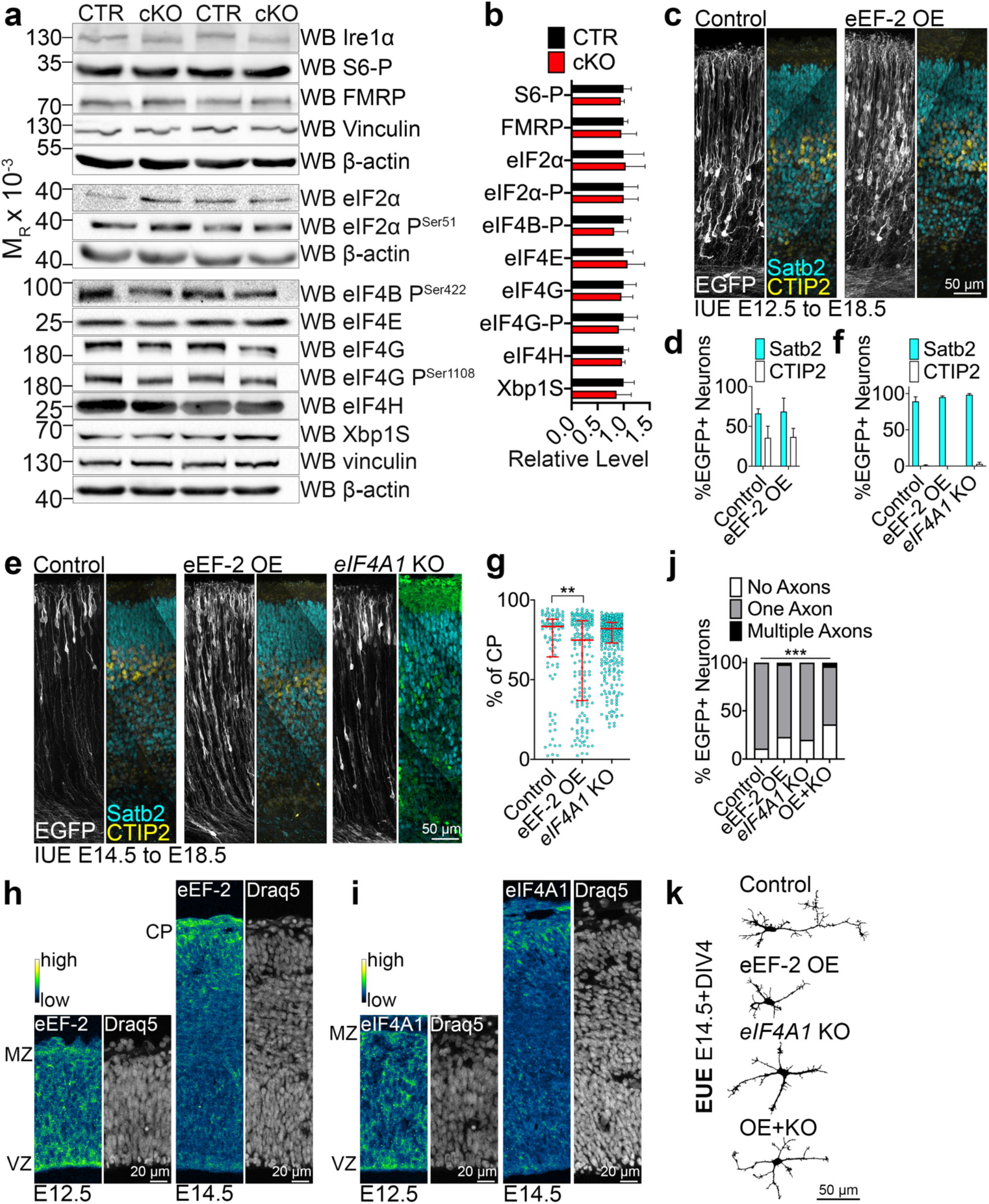
Related to Fig. 3. Ire1α-mediated protein translation regulation employs stress-independent signaling pathways and drives axon specification. (a) Representative results of Western blotting in E18.5 cortical lysates from control and *Ire1α* cKO using indicated antibodies. (b) The expression level of analyzed protein was normalized to the amount of beta-actin in the sample and expressed as relative protein level. (c-g) Representative images (stitched tiles) of EGFP fluorescence signals and immunolabeling for Satb2 and CTIP in E18.5 brains of wild type embryos after IUE at E12.5 (c-d), or E14.5 (e-g) with plasmids encoding for EGFP (Control) or for EGFP and eEF-2 (eEF-2 OE), or for EGFP and eIF4A1 gRNAs and Cas9 nickase (*eIF4A1* KO). Quantification of neuronal identity (d and f) and positioning within the cortical plate (g). (h-i) Representative images of the immunostaining in the coronal cortical slice for indicated proteins and Draq5, a nuclear marker. VZ, ventricular zone; MZ, marginal zone; CP, cortical plate. Graph contains pooled data from indicated number of brains (Table S1). (j) The number of axons projected from a single neuron at DIV4 after EUE to achieve indicated genotypes. Neurons were fixed and immunolabeled from axonal and dendritic markers (compare with Fig. 2). OE+KO, simultaneous eEF-2 OE and eIF4A1 KO. (k) Representative EGFP-based tracings of DIV4 neurons from (j). Apart from Control, shown are neurons with 0 axons. Bar graphs indicate mean ± S.D. Line and error bars on (g) indicate median and interquartile range and scatter plots individual positions of single neurons. For statistics on (b) unpaired t-test with post-hoc Holm-Sidak for multiple comparisons; (d), Mann-Whitney test; (f), (g), Kruskal-Wallis test with Dunn’s multiple comparisons; (j), Chi-square test. 0.001 < ** p < 0.01; *** p < 0.001.

**Fig. S5.**
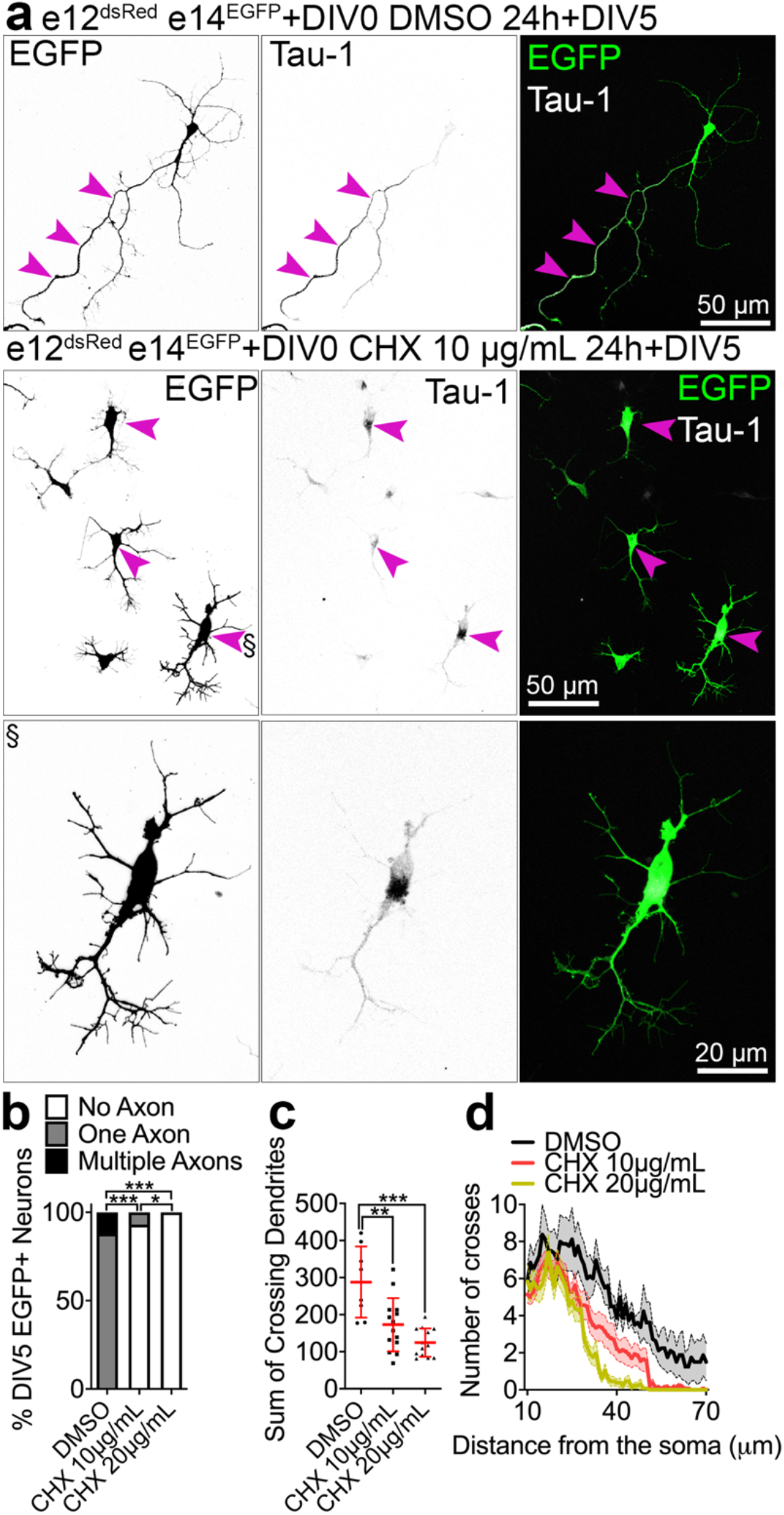
Related to Fig. 7. Axon formation and dendritic growth requires a critical window of translation in precursor cells. (a) Images of immunolabeled primary cells prepared as described in the legend of Fig. 7a. Cells were fixed at DIV5 and immunolabeled for EGFP, dsRed and Tau-1. Purple arrowheads indicate enrichments of Tau-1. The neuron marked with a paragraph symbol is magnified on the bottom panel. (b) Quantification of neuronal polarity. Axon was defined as the neurite with prominent Tau-1 enrichment [compare with top panel in (a)]. (c-d) Quantification of dendritic complexity in DMSO- and CHX-treated DIV5 neurons. Red line and error bars on (c) indicate mean ± S.D. Results on (d) are represented as averages ± S.E.M. For statistical analyses, (b), Chi-square test; (c), D’Agostino and Pearson normality test and one-way ANOVA with Bonferroni post-hoc test. *** p < 0.001; 0.001 < ** p < 0.01; 0.01 < * p < 0.05.

**Fig. S6.**
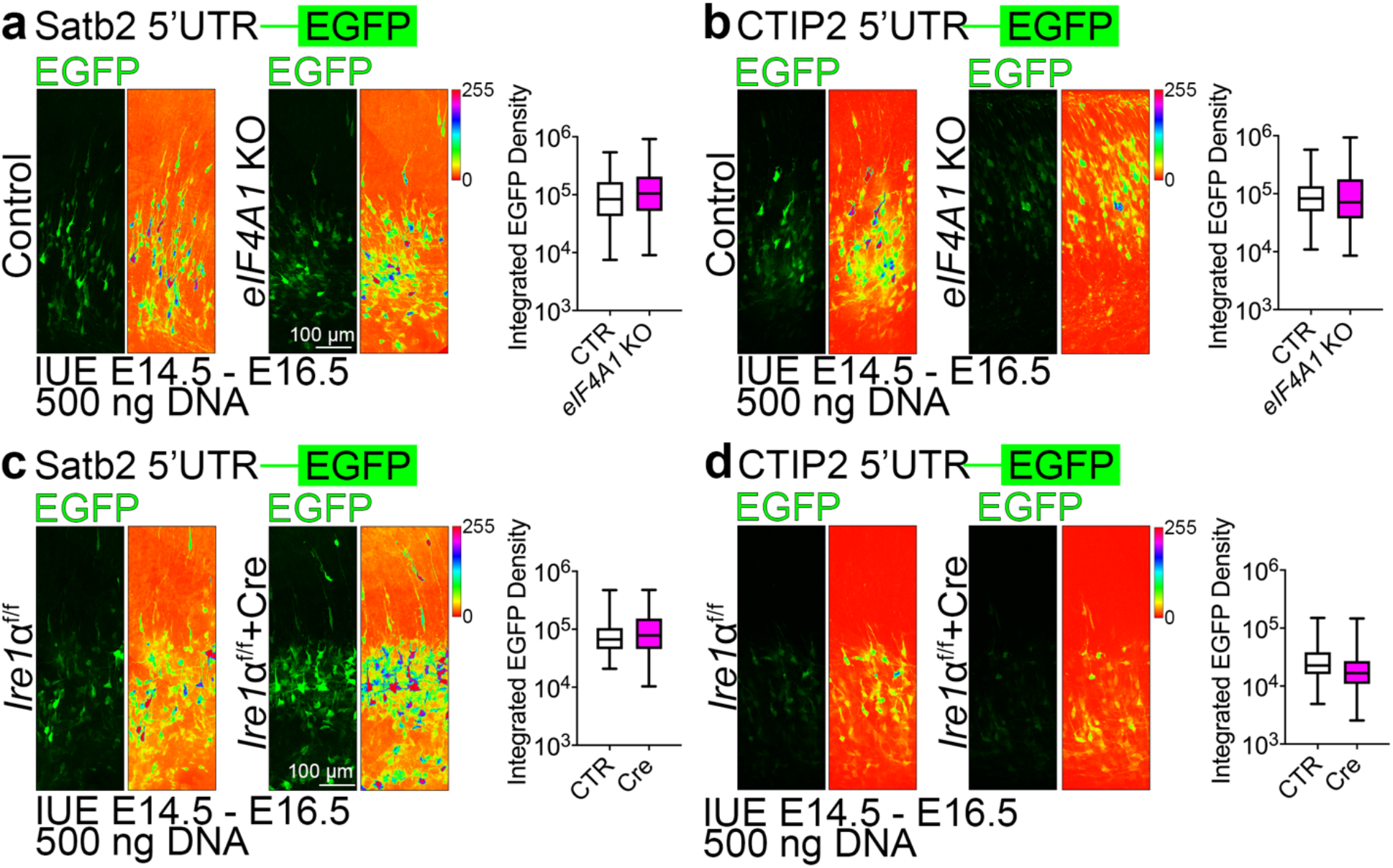
Related to Fig. 8. Inconspicuous eIF4A1- and Ire1α-dependent translation regulation of fate determinants at E14.5. (a-d) Representative images of EGFP fluorescence signals in the E16.5 brain sections after IUE at E14.5 with indicated vectors and Satb2 5’UTR (e) or CTIP2 5’UTR (g) translation reporter construct. Shown are the native signals (left panels) and intensity encoding (right panels). Quantification of EGFP fluorescence signals in single cells expressing translation reporters of Satb2 5’UTR (a, c) or CTIP2 (b, d). Violin plots depict median, interquartile range (box) and minimum and maximum value (whiskers). For statistical analyses, D’Agostino and Pearson normality test and Mann-Whitney test.

**Fig. S7.**
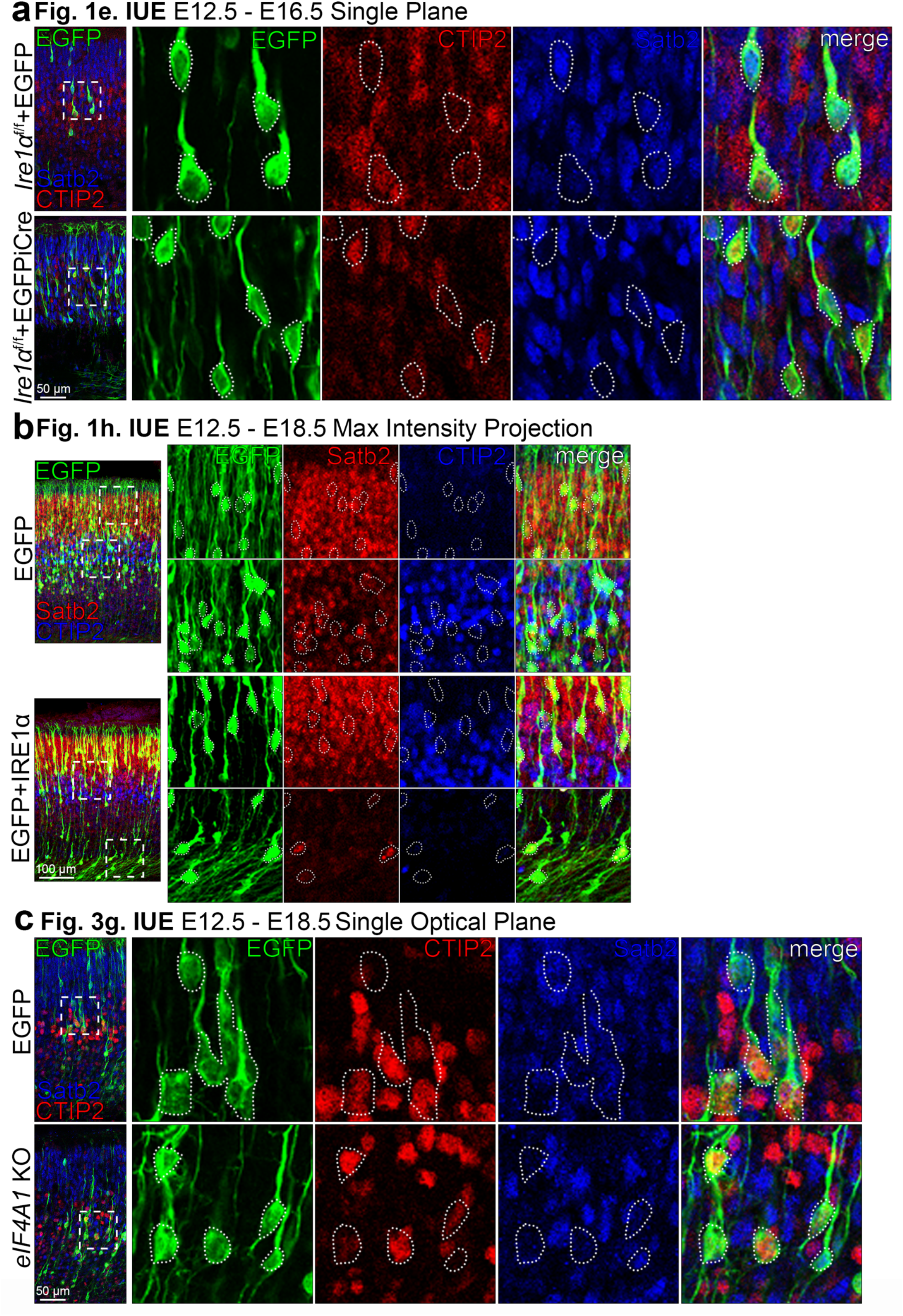
Related to Fig. 1, Fig. 3. Zoom-ins for the neuronal identity phenotypes. (a-c) Representative images of immunostaining against EGFP, Satb2, and CTIP2 in coronal cortical sections after IUE at E12.5 for indicated conditions. Each panel represents a sample image pair used for identity analysis in experiment indicated on top. The squares depict the area of the zoom-in, outlined are somata of electroporated neurons. Neuronal identity was quantified in the proportion of all electroporated neurons in a given coronal section, here shown are snippets for a visualization of immunostaining.

